# GIFT: New method for the genetic analysis of small gene effects involving small sample sizes

**DOI:** 10.1101/2022.02.25.479563

**Authors:** Cyril Rauch, Panagiota Kyratzi, Sarah Blott, Sian Bray, Jonathan Wattis

**Affiliations:** School of Veterinary Medicine and Science, University of Nottingham, College Road, Sutton Bonington, LE12 5RD, UK; School of Life Sciences, University of Nottingham, University Park, NG7 2RD, UK; School of Mathematical Sciences, University of Nottingham, University Park, NG7 2RD, UK

**Keywords:** phenotype-genotype mapping, field theory, complex traits, GWAS, GIFT

## Abstract

Small gene effects involved in complex/omnigenic traits remain costly to analyse using current genome-wide association methods (GWAS) because of the number of individuals required to return meaningful association(s), a.k.a. study power. Inspired by field theory in physics, we provide a different method called Genomic Informational Field Theory (GIFT). In contrast to GWAS, GIFT assumes that the phenotype is measured precisely enough and/or the number of individuals in the population is too small to permit the creation of categories. To extract information, GIFT uses the information contained in the cumulative sums difference of gene microstates between two configurations: (i) when the individuals are taken at random without information on phenotype values, and (ii) when individuals are ranked as a function of their phenotypic value. The difference in the cumulative sum is then attributed to the emergence of phenotypic fields. We demonstrate that GIFT recovers GWAS, that is, Fisher’s theory, when the phenotypic fields are linear (first order). However, unlike GWAS, GIFT demonstrates how the variance of microstate distribution density functions can also be involved in genotype-phenotype associations when the phenotypic fields are quadratic (second order). Using genotype-phenotype simulations based on Fisher’s theory as a toy model, we illustrate the application of the method with a small sample size of 1000 individuals.

## 1. Introduction

Identifying the association between phenotypes and genotypes is the fundamental basis of genetic analyses. In the early days of genetic studies, beginning with Mendel’s work at the end of the 19^th^ century, genotypes were inferred by tracking the inheritance of phenotypes between individuals with known relationships (linkage analysis). In recent years, the development of molecular tools, culminating in high-density genotyping and whole genome sequencing, has enabled DNA variants to be directly identified and phenotypes to be associated with genotypes in large populations of unrelated individuals through association mapping. Genome-wide association studies (GWAS) have become the method of choice, largely replacing linkage analyses, because they are more powerful for mapping complex traits, that is, they can be used to detect smaller gene effects, and they provide a greater mapping precision as they depend on population-level linkage disequilibrium rather than close family relationships. For example, the 2021 NHGRI-EBI GWAS Catalogue currently lists 316,782 associations identified in 5149 publications describing GWAS results (1). Additionally, extensive data collection has been initiated through efforts such as the UK Biobank (2), Generation Scotland (3) and NIH All of Us research program (https://allofus.nih.gov/) with the expectation that large-scale GWAS will elucidate the basis of human health and disease and facilitate precision medicine.

Whilst genomic technologies used to generate data have rapidly advanced within the last 20 years, the statistical models used in GWAS to analyse the data are still predominantly based on Fisher’s method published than 100 years ago (4,5). Using probability density functions (PDFs) and in particular the normal distribution, Fisher’s method partitions genotypic values by performing a linear regression of the phenotype on marker allelic dosage (6). The regression coefficient estimates the average allele effect size, and the regression variance is the additive genetic variance due to the locus (7). Whilst Fisher’s method has been improved, for example using conditional probability linked to potential prior knowledge of genetic systems (Bayes’ method) (8,9), the overall determination of genotype-phenotype mapping is still grounded on PDFs. However, the use of PDFs become problematic in the case of complex/omnigenic traits as they require large scale-study or equivalently, large sample size.

The results obtained by GWAS have demonstrated that complex traits, are driven by a vast number of tiny-effect loci, namely a vast number of genes each with tiny-effect, and not by a handful of moderate-effect loci as initially thought. In turn, this has led to a re-conceptualisation of the genetic basis of complex traits from being polygenic (handful of loci/genes) to omnigenic (vast number of loci/genes) (10–16). Although the omnigenic paradigm is central to further our understanding of biology, there is a practical issue concerning the extraction of information to relate genotype to phenotype in this case. Indeed, tiny-effect loci (i.e., very small gene effects) necessitate a remarkably large population to extract information. Figure 1 exemplifies the limit of GWAS with a restricted sample size of 1000 individuals. This issue regarding the need for large sample sizes was present, but dismissed, in Fisher’s seminal work (4) as he assumed an ‘infinite population’ from the start to use the normal distribution density function in the continuum limit. This assumption allowed him to provide a method able to extract, in theory, the genetic information required to map any genotype to phenotype.

**Figure 1:**
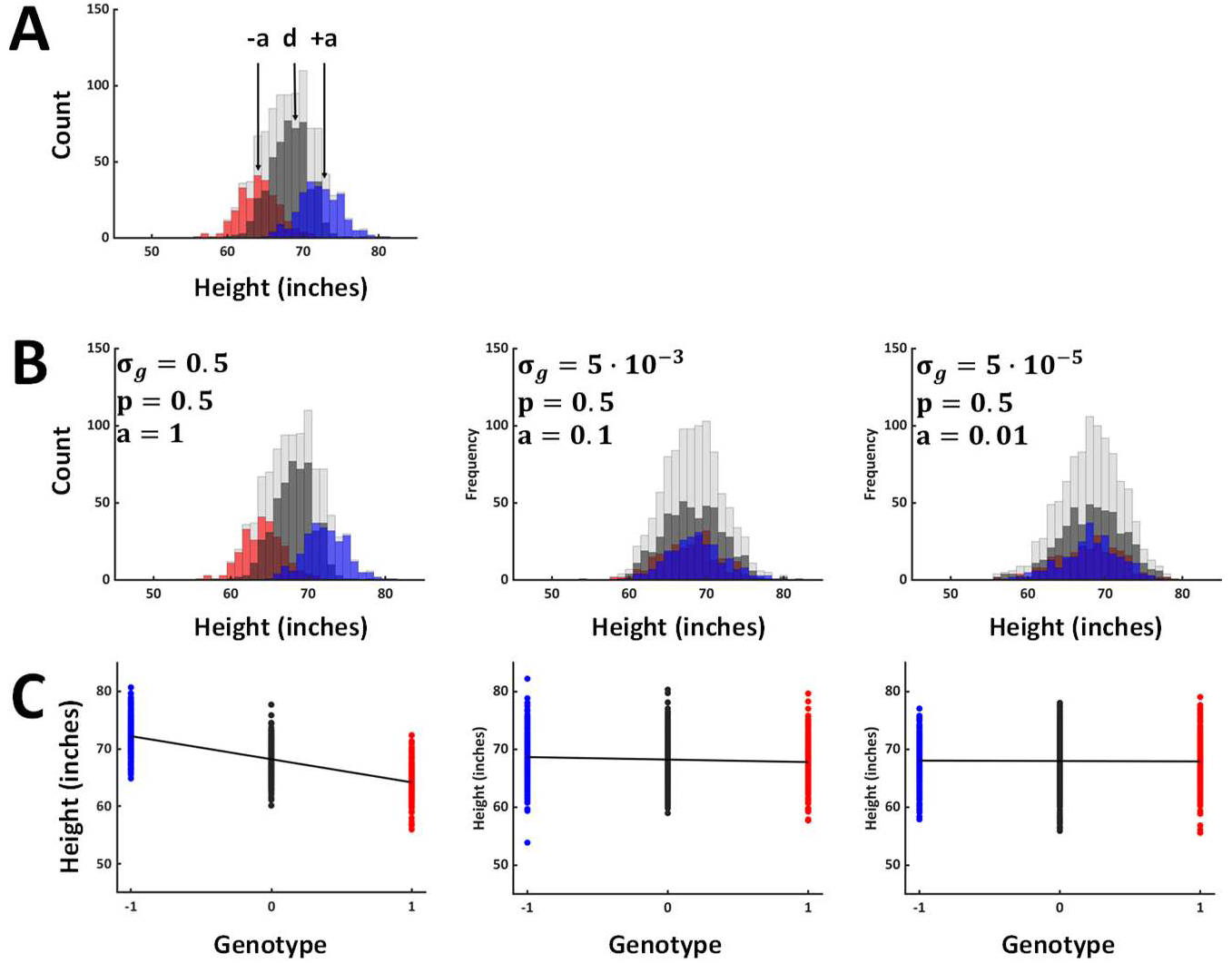
Phenotype human height simulated using the real data from the Genotype-Tissue Expression (GTEx) project (28). We recall that for diploid organisms, such as humans, and for a binary (bi-allelic, A or a) genetic marker, any microstate (genotype) can only take three values that we shall write as ‘+1’, ‘0’ and ‘−1’ corresponding to genotypes aa (red), Aa (grey) and AA (blue), respectively. In the figures the letter, p, refers to the allele frequency and, σ_g_, to the genetic variance. Table 1 provides a relation between the two aforementioned variables. **(A)** Phenotype histograms for the three genotypes (microstates), showing the ‘+1’ state in red, ‘0’ state in dark grey and the ‘−1’ state in blue. The overall phenotype distribution is in light grey at the back. The mean genotypic (microstate) values are −a, d, and +a. In genetics, a and d, are defined as the gene effect and the dominance, respectively. **(B)** Example of limitations linked to current GWA methods when the gene effect, a, changes by one order of magnitude from a=1 to a=0.1 whilst the respective number of microstates does not. In this context, the three populations of genetic microstates collapse, and their separation becomes very difficult to characterise unless the size of the population is increased to reduce the width of categories. In this example, it is almost impossible to distinguish between a=0.1 and a=0.01. Note that in this case the gene effect has been normalised by the phenotypic standard deviation of the population and there is no dominance (d=0). **(C)** Corresponding distributions of the different genetic microstates singled out based on the phenotype values they characterise. The method of averages, or method of moments, advocated by Fisher suggests plotting a straight line as best fit between the three distributions out of which the slope of the straight line is then indicative of genotype-phenotype association. However, the method of averages discards the information that is available in the spreading of data for each genotype. The method we suggest will make use of this information to describe genotype-phenotype association. Note that dominance can occur (d≠ 0) at which point and with the method of average a linear regression of the genotype means, weighted by their frequency, on the number of alleles needs to be performed to provide a new intercept and slope (see (6)).

**Table 1:** Genetic variances as a function of the gene effect and allele frequency.

|  |  | Allele Frequency (p) |  |  |  |  | Genetic variances |
| --- | --- | --- | --- | --- | --- | --- | --- |
|  |  | 0.1 | 0.2 | 0.3 | 0.4 | 0.5 |  |
| Gene effect in unit of phenotypic standard deviations ($a/\sigma$) | 1 | **0.18** | **0.32** | **0.42** | **0.48** | **0.5** | |
|  | 0.8 | **0.1152** | **0.2048** | **0.2688** | **0.3072** | **0.32** |  |
|  | 0.5 | **0.045** | **0.08** | **0.105** | **0.12** | **0.125** |  |
|  | 0.3 | **0.0162** | **0.0288** | **0.0378** | **0.0432** | **0.045** |  |
|  | 0.15 | **0.00405** | **0.0072** | **0.00945** | **0.0108** | **0.01125** |  |
|  | 0.1 | **0.0018** | **0.0032** | **0.0042** | **0.0048** | **0.005** |  |
|  | 0.05 | **0.00045** | **0.0008** | **0.00105** | **0.0012** | **0.00125** |  |
|  | 0.01 | **0.000018** | **0.000032** | **0.000042** | **0.000048** | **0.00005** |  |
|  | 0.005 | **4.5E-06** | **0.000008** | **1.05E-05** | **0.000012** | **1.25E-05** |  |
|  | 0.001 | **1.8E-07** | **3.2E-07** | **4.2E-07** | **4.8E-07** | **5E-07** |  |
|  | 0.0005 | **4.5E-08** | **8E-08** | **1.05E-07** | **1.2E-07** | **1.25E-07** |  |

Whilst one may assume an infinite population mathematically, in practice this comes at a huge cost. To give a ‘real-life’ example of the sample size needed to study complex traits the best is to turn to the phenotype ‘height’ in humans. The phenotype height in humans is a classical quantitative trait that has been studied for over a century as a model for investigating the genetic basis of complex traits (4,17) and whose measured heritability is well known (4,18,19). However, this phenotype has remained controversial (12) for a long time as current association methods were not been able to fully recover the heritability measured (21,22). Whilst different reasons were put forward to explain this discrepancy including, for example, too restricted sample sizes, too stringent statistical tests or the involvement of the environment (6,23); this point seems to have been resolved only recently. Using a population containing a staggering 5.3 million individuals a recent study claims to have captured nearly all of the common single nucleotide polymorphisms (SNPs)-based heritability (24).

This important study confirms that the precision of current quantitative genetic methods to determine omnigenic traits comes at an astronomical cost in line with the assumptions used, namely the need for a staggeringly large (near infinite) population. In this context one may wonder whether such large-scale study will ever be replicated in any other species and in particular those near extinction where small sample sizes need to be considered. Alternatively, one may try to understand where the need for large sample sizes comes from and determine whether it is possible to extract information linking genotype to phenotype in a different way.

There is a very good reason as to why large populations will always be required for omnigenic phenotypes when GWAS is used. As mentioned above, the reason is rooted in the fact that GWAS is mostly based on frequentist probabilities a.k.a. probability density functions (PDFs). Indeed, GWAS is based on statistics and, by definition, statistics deals with the measurement of uncertainties (25). To draw inferences from the comparison of large datasets, a method that requires some understanding of its accuracy, including ways of measuring the uncertainty in data values, is needed. In this context, statistics is the science of collecting, analysing, and interpreting data, while PDFs defined through the notion of relative frequencies, is central to determining the validity of statistical inferences. In practice, the use of frequentist probabilities (or PDFs) and the resulting categorisation of data is justified when inaccuracy exists in experimental measurements. For example, measuring a continuous phenotype such as the height of individuals with a ruler with centimetre graduations, that is, to the nearest centimetre, warrants the use of frequentist probability (or PDFs). In this case, a frequency table of phenotypic values can be defined through 1 cm-width bins or categories, from which the PDFs of the phenotype height and of the genotypes can be deduced to address the statistical inferences. However, the precision available for the inferences will always be, at best, given by the width of categories created and linked to the experimental precision achieved (1 cm in this case). Consequently, if instead of using a ruler with centimetre graduations one was using a ruler with millimetre graduations to increase the precision in inferences, a larger sample size would be required such as to match the new 1 mm-width of categories to reform the PDFs. The trading between the sample size and the precision achieved by GWAS is known as ‘study power’ and its *raison d’être* is linked to the fact that the entire field of probability, and therefore the PDFs, has been conceptualised mathematically to represent the fact that information on a system is limited. It is for this reason that the normal distribution was known before as the ‘error function’ or ‘law of errors’, where the term ‘error’ is defined experimentally (see rulers above). Accordingly, the creation of categories implies that a sort of ‘imprecision’ is necessary.

Whilst the notion of imprecision can be genuine (see rulers above), the act of creating categories to use PDFs when precision in experimental measurements is available is not fully justified and can be seen as an act of ‘wilful ignorance’. This is so because information is lost by slotting different phenotypic values into the same category. To exemplify this point let us take an example, imagine a species close to extinction (very small population, say 50 individuals) and that it is possible to measure phenotypic values with very high precision, for example, using highly advanced imaging techniques or biosensing technologies (26). In this case, each measured individual could return a unique phenotypic value. Consequently, reforming categories to reform and use PDFs would mean embracing the relatively large width of categories created leading to the impression that a large imprecision is present, even so such imprecision did not exist in the first place. One may argue that PDFs, such as the normal distribution in Fisher’s theory, are not required since averages and variances can be mathematically calculated directly from data without the need to recreate PDFs. However, this argument is not valid as averages and variances (and any other moments) cannot be dissociated from PDFs, since they are the ontological parameters that define PDFs, and that PDFs are used to determine statistical inferences in the field of quantitative genetics. Consequently, thinking in term of averages and variances necessitate to conceptualise and hold valid probability density functions, i.e., categories, to describe any system.

Therefore, there is a need to formulate new methods using the full information generated through accurate and highly precise genotyping/phenotyping when sample sizes are small which does not require the categorisation of data. In fact, this problem is equivalent to finding a way to resolve genotype-phenotype mapping by assuming a finite-size population with phenotype values measured precisely enough to rule out the possibility that two phenotype values are in the same category. Taking this challenge as the starting point, a new and relatively simple method for extracting information for genotype-phenotype mapping can be defined. While this method is remarkably simple when explained in lay terms, its theoretical framework requires the introduction of a new concept called ‘phenotypic fields’. Phenotypic fields can also be defined within the context of Fisher’s theory.

The remainder of this paper is organised as follows. In the first part, an intuitive approach to the method GIFT is presented in which one shall see that association between datasets (i.e., genotype and phenotype) can be analysed in specific way that do not involve the use of means and variances necessarily, but phenotypic fields instead. This is followed by a second part stating and explaining the necessary ingredients from physics (entropy, energy and field) and how they must be combined, to provide GIFT. Since GIFT is not too difficult to model, we have relegated the theoretical development of GIFT in the Appendices. Finally (third part) one will demonstrate how Fisher’s seminal theory can be re-transcribed using GIFT. In particular one shall see that Fisher’s seminal intuition corresponds to the simplest form of GIFT. Finally (fourth part), one will compare GIFT to GWAS using simulated genotype and phenotype to demonstrate that GIFT outperforms GWAS.

## 1. Position of the problem and heuristic presentation of GIFT a method

The practical issue regarding genotype-phenotype mappings with current statistical methods concerns the sample size needed to provide accurate/precise information when complex/omnigenic traits are involved. As stated in the introduction, this issue stems from the creation of categories historically linked to the notions of ‘imprecision’ or ‘error’ in measurements. At the dawn of the 21^st^ century we are getting more precise in our measurements, and one may wonder what sort of scientific/mathematical tool we should be using if one were able to attain any level of precision wanted in cases where the population size studied is limited.

We recall here that for diploid organisms, such as humans, and for a binary (bi-allelic, A or a) genetic marker, any microstate (genotype) can only take three values that we shall write as ‘+1’, ‘0’ and ‘−1’ corresponding to genotypes aa (homozygote), Aa (heterozygote) and AA (homozygote), respectively.

One way to proceed to develop a method embracing precision is to start by looking at how density distribution functions are transformed when precision in phenotypic measurements increases. From Fig.2A, the conclusion is obvious, the bar charts are transformed into code bars, where each bar originates from a particular phenotype value representing one individual from the population studied. This result is expected since when the width of categories decreases due to an increase in precision in measurements, there will be a point where there can only be one individual per category. To extract information from the code bars represented by the bottom right chart in Fig.2A, let us now wonder what it means to have information on the phenotype as opposed to have none.

**Figure 2:**
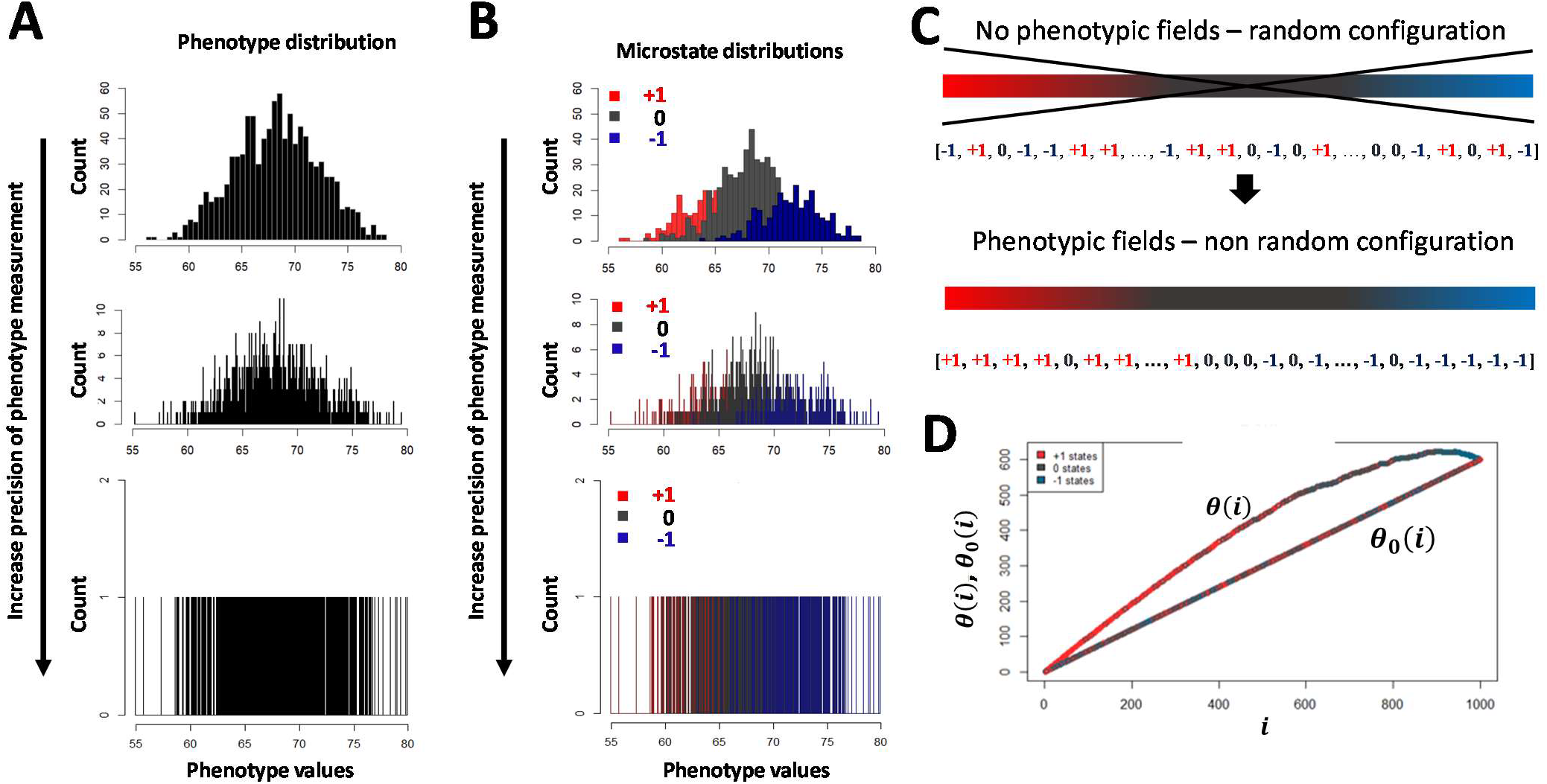
**(A) & (B)** When applied to real data sets, current genome-wide association studies rely on probability distribution density functions (PDFs) namely the creation of frequency plots (method of relative frequencies) via the grouping of phenotype values into categories representing range of phenotype values (A, top-chart). The same method (PDFs) is then applied to genotypes (B, top chart). We recall that for diploid organisms, such as humans, and for a binary (bi-allelic, A or a) genetic marker, any microstate (genotype) can only take three values that we shall write as ‘+1’, ‘0’ and ‘−1’ corresponding to genotypes aa (red), Aa (grey) and AA (blue), respectively. The comparison of the two top charts in (A) and (B) demonstrates how genotype are associated with the phenotype, as in this case any phenotype category can be decomposed using the underlying microstate categories. However, grouping data into categories is legitimate so long that the width of the category is justified. The width of categories is justified provided that imprecision exists in phenotype measurements. For example, if height in human was the phenotype of interest measured with a ruler with inch graduations, namely measured to the nearest inch (scale of imprecision), then the width of categories would be 1 inch. However, a method based on the notion of imprecision has limited value when precision is available, and new methods are required in this case. Indeed, by increasing the precision in phenotype measurements it is possible to envision, in a near future, the possibility to deal with genotype and phenotype under the form of ‘code bars’ (A & B, bottom-charts) as opposed to PDFs. The question is then, how can information be extracted from those ‘code bars’? **(C)** To analyse the ‘code bar’ in the bottom charts of (A&B), we rewrite it as a string of microstates. The first thing to note is that the way the microstates ‘+1’, ‘0’ and ‘−1’ appear in the string (a.k.a. configuration/ordering), is linked to the fact that: (i) the genome position considered is associated with the phenotype and, (ii) phenotypic values are ranked as a function of their magnitude. Accordingly, if the genome position considered is not associated with the phenotype, the configuration would appear random (top string) as opposed to being ordered if a genotype-phenotype assoxaition exist (bottom string). **(D)** Thus, to determine as to whether a genome position is associated with the phenotype, the best way is to plot the cumulative sum of microstates and to compare it to the plot of its ‘scrambled’/random configuration. When the string is ordered, a curve will emerge as shown by *θ*(*i*). On the contrary, when string is ‘scrambled’ a straight line is expected as shown by *θ*_0_(*i*). The straight line resulting from scrambling of the string is expected as the presence probabilities of pulling ‘+1’, ‘0’ and ‘−1’ from the scrambled string are constants in this case. The curve (ordered string) can be modelled by considering that the information about the phenotype acts like a ‘field’ to change the configuration of microstates from being disorderd to being ordered. The two curves intersect at the last position in the string because the numbers of ‘+1’, ‘0’ and ‘−1’ in either string (ordered or scrambled) is conserved. The difference between the ordered and disordered strings provides a precise measure of the genetic influence on the phenotype. In Figure 4A we provide a plot of such a difference in the phenotypic space.

To answer this question the best thing is to further simplify the problem by considering the coloured bars only and not their spacing. Imagine, therefore, that a set of individuals has been genotyped and that those individuals are picked at random. That is, there is no information on any phenotype. Imagine also that one decides to concentrate, for example, on the genome position 1’000’000 on chromosome 4 for all the individuals since this genome position happens to display a biallelic single nucleotide polymorphisms (SNPs) across the set of individuals.

Thus, upon calling randomly but sequentially individuals, the genotypic information obtained in due course can be represented as a random string of genotypes including ‘+1’, ‘0’ and ‘−1’ microstates (representing homozygote-AA, heterozygote-Aa and homozygote-aa). An example of such random configuration is:

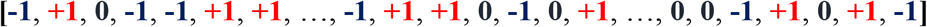

Note that the order in which the individuals were called is linked to the position in the string. Let us now repeat the same experiment using the same individuals in a context where accurate information on a chosen phenotype is available. That is, we call the individuals as a function of the magnitude of their phenotype we consider. For example, if the phenotype is height, one starts by calling the smallest individual and all subsequent individuals through successive increments in their phenotype height. Note again that because each individual has a unique phenotype value there is no possibility for two individuals to be called at once.

If the genome position 1’000’000 on chromosome 4 is involved in the formation of the phenotype, then one would expect a change in the configuration of the string of microstates based on the fact that homozygotes would be found at the extremities of the string and heterozygotes towards the middle (see Figure 2). An example of such a string would be, for example:

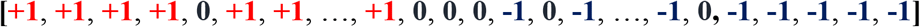

Thus, the only thing that changes between the random and the phenotype-ordered configurations is the way the genetic microstates are allocated to positions in the string. However, as the genome position 1’000’000 on chromosome 4 is the only one that has been considered, the two configurations contain the same number of ‘+1’, ‘0’ and ‘−1’, since the same individuals were considered between the two configurations.

The *ansatz* is then to consider the cumulative sum of microstates as a function of the position in the string. Indeed, it is clear from the examples given above that if one starts by adding the microstates together, differences will be seen in the resulting cumulative sums. To give an example, let us consider the two strings above and note ‘*θ*_0_(*i*)’ and ‘*θ*(*i*)’ the cumulative sums of microstates in the random and ordered configurations, respectively, where ‘*i*’ is the position in the string. Then adding the microstates starting from the left side of the strings one finds:

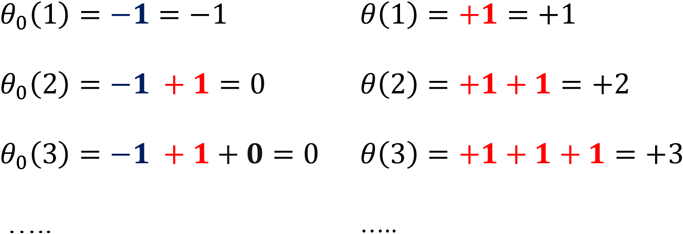

As a result, the difference ‘*θ*(*i*) − *θ*_0_(*i*)’ is expected to be indicative of the importance of the phenotypic information and how gene microstates are related to the phenotype. The fact that the same individuals were considered in both configurations also impose a conservation relation under the form: *θ*(*N*) − *θ*_0_(*N*) = 0. One shall call the cumulative sums: ‘genetic paths’ whose mathematical definition will be précised below. To conclude, it is the information on the phenotypic values that provides a change in the configuration of microstates and one can start developing the formulations of, *θ*_0_(*i*) and *θ*(*i*).

Noting, N_+1_, N_0_ and N_−1_ the number of genetic microstates ‘+1’, ‘0’ and ‘−1’, respectively. The genetic microstate frequencies for genome position 1’000’000 on chromosome 4 are defined by, 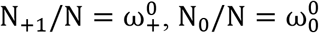 and 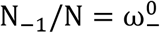.

When the positioning of the genetic microstates in the string is performed in a random fashion, the probabilities of finding ‘+1’, ‘0’ or ‘−1’ as genetic microstate at any position ‘i’ are 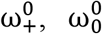 and 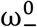, respectively. The resulting cumulative sum is then: 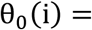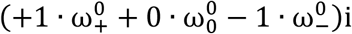. Consequently, θ_0_(i) is therefore a straight line. We shall call θ_0_(i) the ‘default genetic path’(Fig.2C).

In the second configuration the microstates are ordered. Noting ω_+_(i), ω_0_(i) and ω_−_(i) the occurrence probabilities of the genetic microstates ‘+1’, ‘0’ and ‘−1’ the cumulative sum is then: 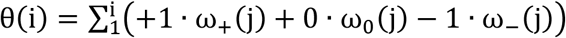; where θ(i) is defined as the ‘phenotype-responding genetic path’ (Fig.2C).

As a result, the signature of a gene interacting with the phenotype when considering the two aforementioned genetic paths is the difference: 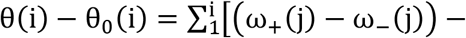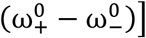. One can then be a little bit more prescriptive by introducing the notion of phenotypic fields.

## 2. A Physics-inspired model for GIFT: Notion of ‘phenotypic fields’ and resulting difference between the phenotype-responding and default genetic paths

The difference, θ(i) − θ_0_(i), can be described using field theory. Indeed, as the only difference between the two configurations is the information linked to the phenotypic values, the phenotypic information can be thought as an external field impacting the configuration of microstates. To provide a physics-inspired definition of genotype-phenotype mapping, let us reconsider the random string above and assume that the set of individuals in the string are particles and that the different genetic microstates ‘+1’, ‘0’ and ‘−1’ are their physical properties. One can then assume that it is those properties that interact with the field. Note that contrary to physics where a single field is defined, one needs in our case to define one field per microstate. Namely that the ‘field’ aforementioned is, in fact, composed of sub-fields. One shall note by u_+_(Ω), u_0_(Ω) and u_−_(Ω) the phenotypic fields acting on the microstates ‘+1’, ‘0’ and ‘−1’ respectively. Note that the variable Ω represents the phenotypic values measured precisely. By assuming further that the particles cannot interact together and that, when they are not forced into a specific configuration by the fields, they can hop and exchange positions when the field is null (similar to a diffusion/thermal process), one can then model the string of microstates as a closed system. Fig.2C provides an idea of how the ‘phenotypic fields’, when non null, impacts on the configuration of microstates. With those assumptions and using basic principles from statistical physics it is then possible to model the presence probability of microstates at any position in the string.

Thus, after re-expressing the genetic paths in the space of phenotypic values since the fields are function of the phenotypic values (Appendix 1), one can then construct functionals representing the entropy (Appendix 2) and the total interaction energy between the microstates and the subfields (Appendix 3). Finally, one can optimise a functional similar to the free energy to express how the fields are related to the asymmetry of states (Appendix 4). Consequently, one can demonstrate the familiar result concerning the presence probability of microstates expressed in the phenotypic space as,

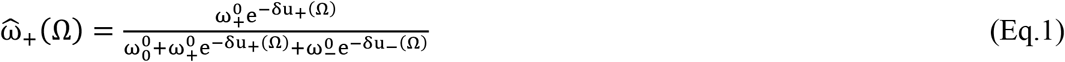

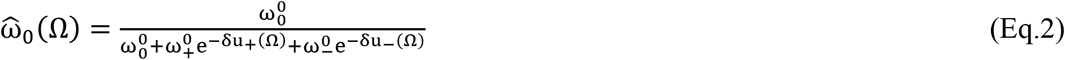

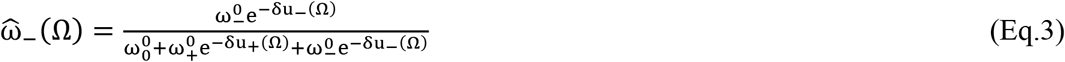

Where the hat 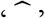 is added to insist on the fact that the presence probabilities of microstates are expressed in the space of phenotypic values (and not positions) and, 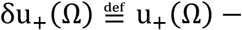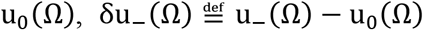. Eqs.1–3 are familiar to physicists when dealing with Boltzmann’s weigh in statistical physics. Note that the default genetic path is defined when the fields are null. Noting, 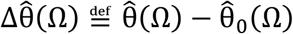, the difference between the phenotype responding and default genetic paths expressed in the phenotypic space, 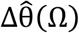 is therefore a function of the difference between Eq.1 and Eq3. One can then make the symmetries of the problem more apparent by defining for the genetic microstates, 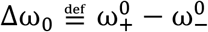 and 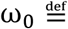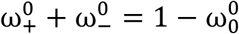; and for the phenotypic fields, 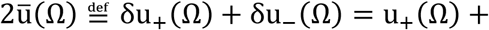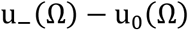 and 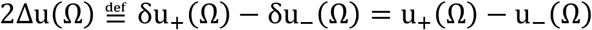. In this case, using hyperbolic functions one deduces (see Appendix 4 for development),

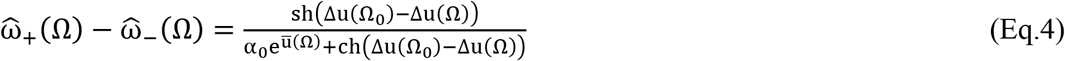

Where, 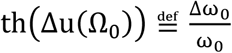 and 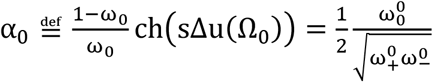. The new variable ‘Ω_0_’ is the phenotype value corresponding to the condition 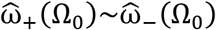 and the meaning of the constant ‘α_0_’ can be related to the Hardy-Weinberg law from population genetic. Hardy-Weinberg law based on random mating in a population provides a relationship between the genetic microstate frequencies under the form: p^2^ + 2pq + q^2^ = 1, where p^2^ and q^2^ are the genotype frequencies of genetic microstates ‘+1’ and ‘−1’, i.e. homozygote genotypes aa and AA, respectively; and 2pq the genotype frequency for genetic microstate ‘0’, i.e. the heterozygote genotype Aa. In our case, this corresponds to replacing p^2^, q^2^ and 2pq with, respectively, 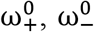 and 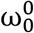. Consequently, the Hardy-Weinberg law imposes α_0_ = 1 with α_0_ ≠ 1 corresponding to a deviation from the law. However, this term is expected to remain stable upon any changes of allele or genotype frequencies suggesting therefore that, genetically, any changes in ‘Δω_0_’ are to some extent compensated by corresponding changes in ‘ω_0_’. Finally, using Eq.4 one deduces the difference between the phenotype responding and default genetic paths expressed in the phenotypic space,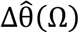, as (Appendix 5):

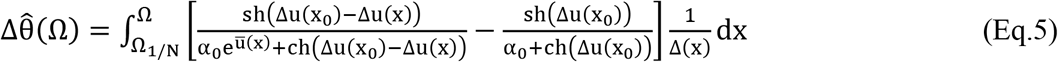

Where Ω_l/N_ is the smallest phenotypic value measured and, Δ(x), is the spacing between individuals in the code bar Fig.2A that can be related to the probability density function of the phenotype when the population measured is dense (see Appendix 1 and SM1 in the Supplementary Materials). Finally, the conservation of genetic microstates needs to be added, that is, 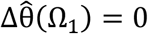, expressed as,

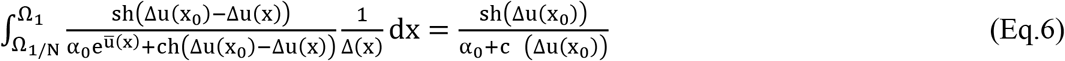

Where Ω_1_ is the largest phenotypic value measured. Therefore, as α_0_ is constant since a single genome position is considered, the genetic paths difference can be re-expressed integrally using two independent reduced phenotypic fields, i.e., Δu(Ω) and 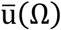, and Eq.6 provides a coupling between those fields and Δ(Ω). The advantage of using fields is the reduction of unknown parameters involved in the problem and the possibility of laying out genotype-phenotype associations based on fields’ symmetry. For example, and as a minimalist model, one may wonder what sort of expression would take the fields if the reference field u_0_(Ω) was null and, sδu_+_ and sδu_−_, were acting anti-symmetrically and linearly on the microstates ‘+1’ and ‘−1’? This minimalist model can be developed (see SM2 in the Supplementary Materials) and is similar to Fisher’s seminal intuition concerning genotype-phenotype associations (see below).

Our aim is now to demonstrate that the idea of genetic paths mediated by phenotypic fields already exists in Fisher theory. This can be shown by coarse graining the paths.

## 3. Coarse graining GIFT: Definition of fields and Fisher’s context, implication for small gene effects and definition of variance fields

### 3.1. Coarse graining GIFT

To derive a coarse-grained version of GIFT, that is, a genetic path difference for GWAS, we assume the existence of categories or bins and concentrate on the interval of phenotype values ranging between Ω and Ω + δΩ defining one particular category or bin.

Based on frequentist probability, by noting by N the total number of individuals in the population we can define by: δN~N ∙ P_Ω_(Ω)δΩ, the number of individuals in the phenotype category concerned namely with a phenotype value ranging between Ω and Ω + δΩ.

Similarly, concentrating on a single genome position, we can define by: 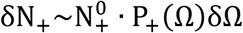, 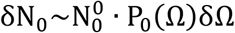 and 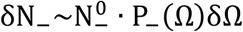 the respective number of ‘+1’, ‘0’ and ‘−1’ genetic microstates for the phenotype values ranging between Ω and Ω + δΩ, where 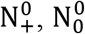 and 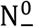 correspond to the total number of ‘+1’, ‘0’ and ‘−1’ microstates in the population with respective presence probability density functions, P_+_(Ω), P_0_(Ω) and P_−_(Ω).

The design of categories generates two conservation relationships: The first one concerns the total number of individuals and microstates, namely, that for a given genome position, the sum of all possible microstates is also the sum of all individuals. This relationship is written as follows: 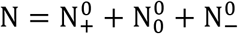. The second conservation relationship is linked to the category considered. The number of individuals in the category concerned is also the sum of the microstates in this category: δN = δN_+_ + δN_0_ + δN_−_. Consequently, the conservation relation concerning the number of individuals and microstates in the concerned category can be rewritten as,

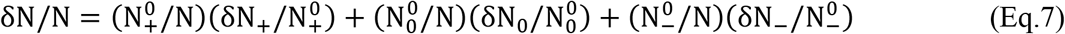

Using the probability density function of both microstates and the phenotype defined above, the following is deduced:

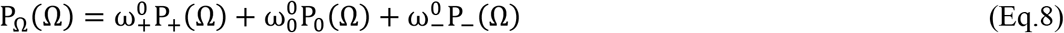

From Eq.8 all the moments of microstate distributions are related to those of the phenotype distribution. Let us note by, 〈Ω〉, and, a_+_, a_0_, a_−_, the average values of the phenotype and microstates ‘+1’, ‘0’, ‘−1’ distribution functions, respectively; and by σ^2^ and 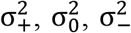 the variances of the phenotype and microstates ‘+1’, ‘0’, ‘−1’ distribution functions, respectively. From Eq.8, one deduces the conservation relations for the first two moments in the form:

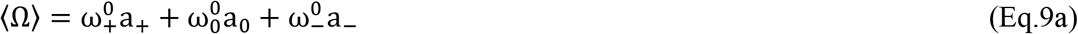

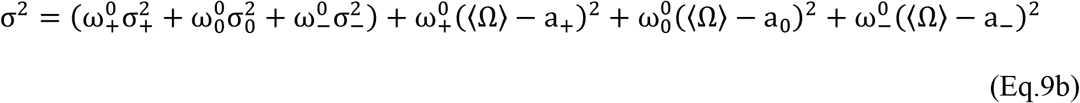

The relations provided by Eqs.9 are valid by definition, namely, whatever probability density functions are involved. Whilst Fisher never formulated a conservation similar to the second from Eq.9b, in his seminal paper (4) he used the notation α^2^ to denote the genetic variance in the form: 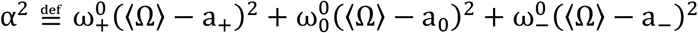.

Let us now define the coarse-grained version of Eq.4 by noting, 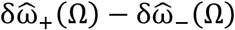, the difference in the presence probability of microstates ‘+1’ and ‘−1’ for the category of interest, it is then deduced:

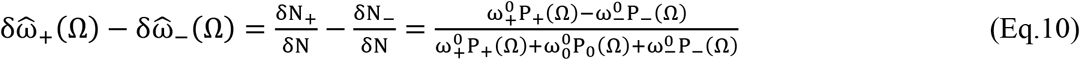

Note that Eq.10 corresponds to a local relative difference, in the space of phenotypic values, of microstates ‘+1’ and ‘−1’. Also, it can be verified that the conservation of gene microstates holds by summing, δN_+_ − δN_−_, over all existing categories or bins, namely: 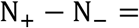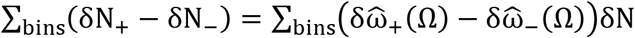. Using the definition δN = NP_Ω_(Ω)δΩ together with Eq.8, we deduce using the continuum limit, 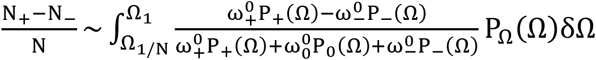 or equivalently, 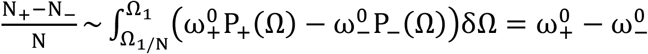.

Direct mapping of fields can then be performed between GWAS and GIFT (SM3 in the Supplementary Materials). As a result, we can define the coarse-grained versions of the difference in the genetic paths using the continuum limit as:

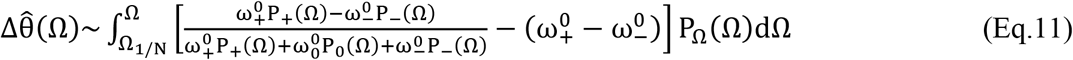

Eq.11 demonstrates that 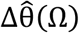 is sensitive to the probability density functions involved as a whole and not just to the average values. In other words, the variance of microstates and their average values will impact on genotype-phenotype associations. Note that in Eq.11 the integration interval is unchanged. However, the convergence in probability of distributions allows some freedom, for example changing the integration interval from [Ω_l/N_; Ω_1_] to [0; +∞[, or from [Ω_l/N_ − Ω_m_; Ω_1_ − Ω_m_] to]−∞; +∞[, where ‘Ω_m_’ is a median position.

The fields can then be defined setting: 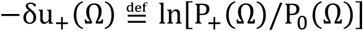 and 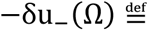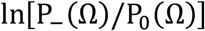; and from those relations it is deduced: 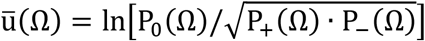 and 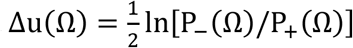.

Finally using Eq.5’s notations one deduces:

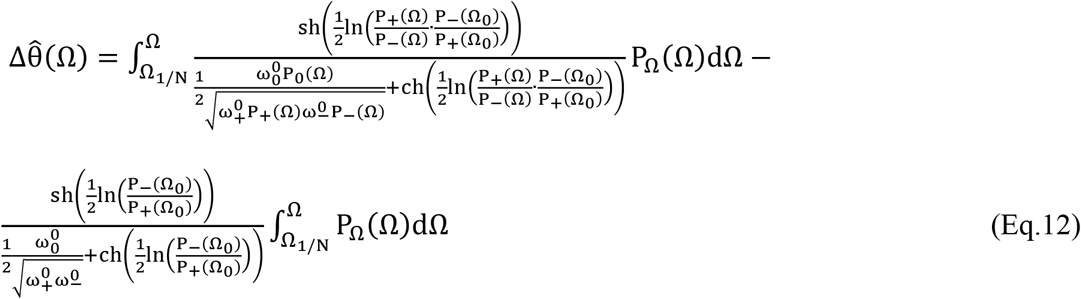

The significance of these fields can now be addressed. The field 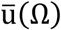 describes local deviations from the Hardy-Weinberg law, valid for each bin or category of phenotype values. For example, if a population was under no selection and random mating occurred, then the whole population would follow the Hardy-Weinberg equilibrium law, that is, 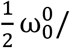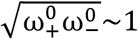. However, selecting a particular bin or category of phenotype values would demonstrate a local deviation of this law, given by the term 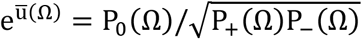. The signification of Δu(Ω) can be addressed using Fisher’s approach.

### 3.2. Definition of fields using Fisher’s theory

In his seminal paper (5), Fisher hypothesised that in a context where the population is infinite to use the normal distribution, the genetic variance ‘α^2^’ is much smaller than the phenotype variance and that the variances of microstate distribution density functions for each gene are similar to that of the variance of the phenotype. Whilst his hypothesis can be understood intuitively when all distribution density functions nearly overlap, it can also be demonstrated using Eq.9b. Indeed, assuming α^2^ ≪ σ^2^ implies σ^2^ − α^2^~σ^2^ and therefore 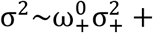 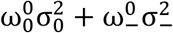. As 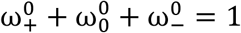, posing 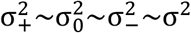 as Fisher did, is one valid solution. However, the relation 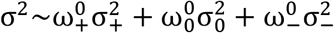, is the equation of an ellipse, and an infinite number of solutions are, in theory, possible. Those solutions will dependent on the variances of microstates (see Section 3.4. below, that is, the definition of fields linked to the variance of microstates).

Following Fisher’s assumption, the probability of finding the microstates ‘+1’, ‘0’ and ‘−1’ as a function of phenotype values are expressed as (4) (Figure 1) : 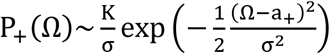, 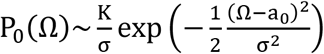 and 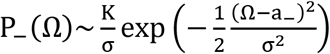. Where ‘K’ is the normalisation constant identical for all microstates distribution functions since they nearly overlap. Noting, 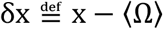, the difference between the variable, x, and the average of phenotype values to simplify the notations, as the second orders defined by (δa_+_/σ)^2^, (δa_0_/σ)^2^ and (δa_−_/σ)^2^ are neglected in Fisher’s context, one deduces then:

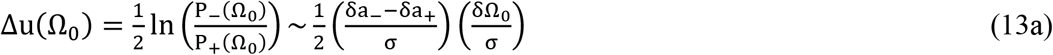

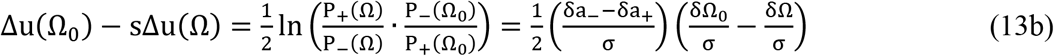

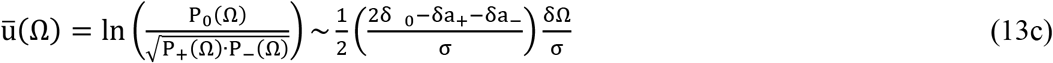

In Eq.13a and Eq.13b the term δa_−_ − δa_+_ = a_−_ − a_+_ = 2a (see Fig.1A) is known as the ‘gene effect’ in GWAS. In Eq.12c, the term 2δa_0_ − δa_+_ − δa_−_ = 2a_0_ − a_+_ − a_−_ = d (see Fig.1A) is the dominance as defined in the GWAS. As it turns out, Fisher considered: d~0.

Altogether, these results demonstrate that Fisher’s theory can be described by phenotypic Fields and genetic paths. As it turns out Fisher’s model corresponds to the minimalist model aforementioned (SM2 in the Supplementary Materials). Using these fields, it is also possible to determine a generic solution to Eq.8 (see SM4 in the Supplementary Materials).

### 3.3. Implication for small gene effects

Complex traits involve genes with very small effects that are difficult to characterise. The aim is to determine the resulting difference in the genetic paths in this case, that is, when the gene effect, a = (a_−_ − a_+_)/2 (see definition above), tends towards zero: a → 0. Because probability density functions are used, the integration interval can be altered using the convergence property of the distributions. In this context, the conservation of genetic microstates (Eq.6) can be written using Fisher’s fields as,

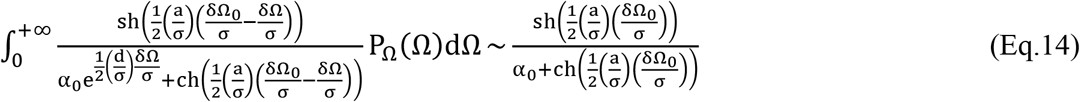

Consider, as Fisher did in his seminal paper (4), a phenotype distribution of the form, 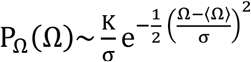, and rescale the phenotype values in the integral using, a/σ, as a scaling parameter as follows:

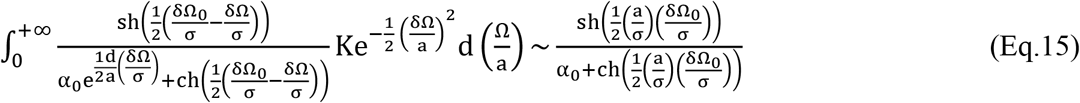

By taking the limit a → 0, the rescaled phenotype distribution becomes a Dirac distribution, dominating any convergences; thus, the left-hand side can be transformed as:

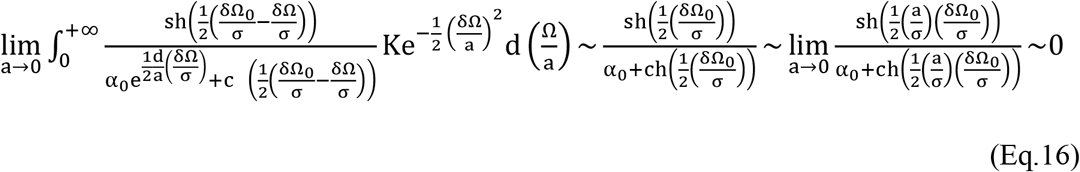

Therefore, small gene effects imply: δΩ_0_/σ ≪ 1. Recalling that 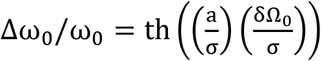, one also deduces 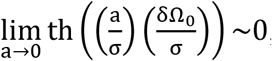, that is, small gene effects always involve common allele frequencies, namely Δω_0_/ω_0_ → 0. Using this result, the genetic paths difference can then be developed when gene effects are small and by assuming that P_Ω_(Ω_l/N_) ≪ 1 and that ‘P_Ω_(Ω)’ is normally distributed one obtains at the leading order:

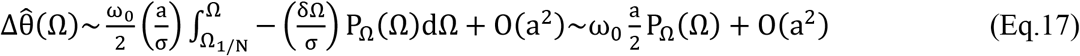

Eq.17 shows that in the context of Fisher’s theory, a small gene effect corresponds to an overlapping symmetry between the genetic microstates and the phenotype distribution, with an amplitude proportional to the gene effect.

### 3.4. Fields linked to the variance of microstates

The involvement of variances in microstate distribution functions in genotype-phenotype associations is a highly debated matter (see (20) and references within). As mentioned above, the expression of the difference of the genetic paths considers the distribution density function as a whole, including the role of the microstate variances. In this context, we saw that Eq.9b provides a relation between variances in the form of an ellipse. Assuming a single variance for all microstate and phenotype distributions, as Fisher did, is plausible; but other solutions exist that would not violate Eq.9b. In this context, let us imagine that the gene effect and dominance are nulls but that the distribution density function of microstates ‘+1’, ‘0’, ‘−1’ and of the phenotype have distinct variances; written, respectively, as: 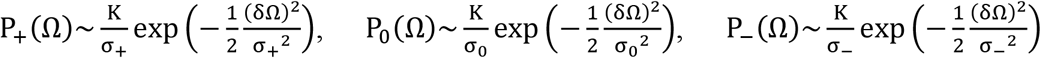 and 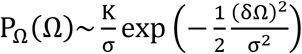.

By noting λ_+_ = σ/σ_+_, λ_0_ = σ/σ_0_ and λ_−_ = σ/σ_−_, the fields can be mapped under the form:

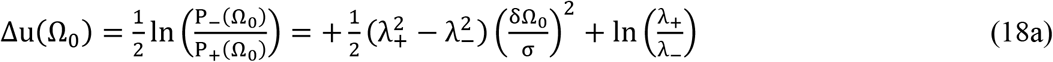

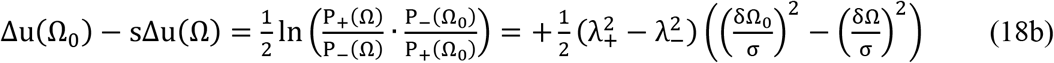

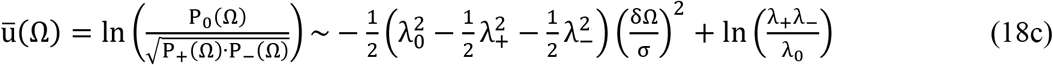

Consequently, pseudo-gene effect and pseudo-dominance linked to the variances of genetic microstates can be defined, respectively, as: 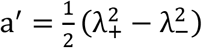 and 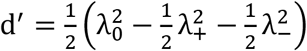.

To conclude, as this new method does not only concentrate on average values, it captures more information as far as genotype-phenotype associations are involved.

## 4. Illustration of the application of GIFT using simulated data

We intend to illustrate how GIFT can be applied using simulated data and qualitatively assess its sensitivity to extract information.

### 4.1. Data simulations

The codes used are provided in SM5, see the Supplementary Materials.

Data was simulated according to quantitative genetic models defined by Falconer and Mackay (1996) (27). A single bi-allelic quantitative trait locus (QTL) associated with a continuous phenotype was modelled, with an additive allele effect, a, and allele frequencies, p and q, where p + q = 1. The simulation parameters were set as the number of individuals sampled, N = 1000; number of simulation replicates, n = 1000; allele frequency, p; additive allele effect, a, and dominance, d.; note that the number of simulation replicates allows one to determine the best outcomes. While the theory provided in this paper is general, the simulation of data will be restricted to individuals’ genotypes allocated according to Hardy-Weinberg proportions. For N individuals, Np^2^ had genotype AA (corresponding to microstate −1), 2pqN had genotype Aa (microstate 0), and Nq^2^ had genotype aa (microstate +1). The allele effect, a, is defined as half the difference between the +1 and −1 genotype (microstate) means, and d, is the position of the 0 genotype (microstate) mean (Figure 1). Dominance is measured as the deviation of the mean of microstate 0 from the midpoint between the means of the +1 and −1 microstates. For the purposes of the simulation dominance, d, was 0, that is, the mean of microstate 0 was mid-way between the mean of microstates +1 and −1.

The additive genetic variance due to the quantitative trait loci 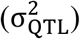 was defined as (27): 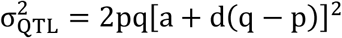. Each individual was assigned a genotypic value, depending on their microstate: −a for the +1 microstate, 0 for the 0 microstate, and +a for the −1 microstate. Individual phenotypes were generated by adding a random environmental effect to the genotypic value of each individual. The added environmental effect was a random variate drawn from a normal distribution with a mean of 0 and variance of 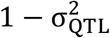. The phenotype was then rescaled to a value representing a realistic dataset: Phenotype = (simulated phenotype × standard deviation of real data) + mean of real data. In this case, the real dataset modelled was a summary of the Genotype-Tissue Expression (GTEx) project (28). In particular, the phenotype was height with a mean of 68.17206 inches and standard deviation of 4.03007 inches. For each simulated replicate of N individuals, the difference between the cumulative sums of microstates ordered by phenotype value and genotypes in a randomised order with respect to phenotype was determined to create the difference in the genetic paths difference. The maximum value of this difference was identified and its position and phenotypic value in the ordered string of microstates were recorded. Where the maximum value extended over several positions, the mean position and phenotypic value were recorded. Finally, to simplify representation, the amplitudes of the genetic path differences were normalised by population size (N = 1000 in this case).

Note that the standard deviation(s) arising from genotype-phenotype simulations were not considered in the analysis that follows. Instead, we report a theoretical analysis of the convergence of the genetic path difference method, and its self-consistency, as well as its sensitivity to detect genotype-phenotype associations using simulations, in SM6 and SM7, respectively, see Supplementary Materials.

### 4.2. Analysis of simulated results

For information, Table 1 shows how genetic variance, gene effect, i.e., a/σ, and allele frequency are numerically related using GWAS method. Similarly Figures 3A and 3B represent, in the context of GWAS and for the allele frequencies p=0.5 (Δω_0_ = 0) and p=0.8 (Δω_0_ = 0.6) that will be used below as examples, the relationship between the power of the study, the gene effect and the sample size as described in (29). Briefly, the power of a study is related to the concepts of Type I and Type II errors. A Type I Error (a.k.a. α) is rejecting the null hypothesis in favor of a false alternative hypothesis, and a Type II Error (a.k.a. β) is failing to reject a false null hypothesis in favor of a true alternative hypothesis. The power of a study is then the probability of avoiding a Type II error. Mathematically, the power is defined by, 1 – β, where 0<β<1. If the power is close to 1, i.e., β~0, the hypothesis test is very good at detecting a false null hypothesis. β is commonly set at 0.2, to provide a power ~0.8 (or 80%). Powers lower than 0.8, while not impossible, would typically be considered too low for GWAS. The four primary factors affecting power are, the sample size, the significance level (or α), the variance/variability in the measured response and the magnitude of the effect of the variable. Only the first variable can be altered in a study since all the others are fixed by the genes. To conclude, power is increased when the sample size or effect sizes (gene effect) are increased. Accordingly, Figure 3 demonstrates that 1000 individuals would not allow 80% power to be achieved unless the gene effect is sufficiently large, that is for a/σ ≥ 0.5.

**Figure 3:**
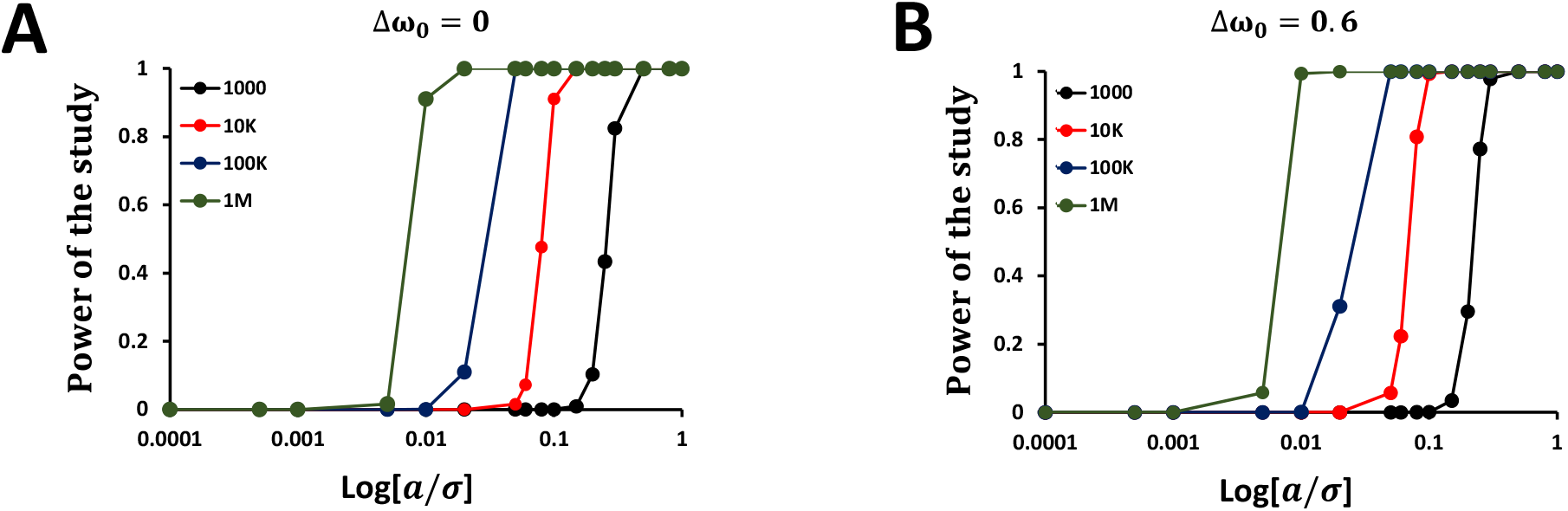
Power calculations for two allele frequencies as a function of the gene effect and sample size using Fisher’s additive model when the dominance is null. The colour of the curves is associated with the sample size (one thousand individuals for black, ten thousand individuals for red, hundred thousand individuals for blue and one million individuals for green). The figure illustrates that with 1000 individuals current genotype-phenotype methods are not powerful enough to detect small gene effects.

Using simulated data, we can now represent the genetic paths difference and its log transformation for Δω_0_ = 0, either in the space of positions (Figure 4A) or of phenotype values (Figure 4B) for different log-scale values of the normalised gene effect a/σ.

**Figure 4:**
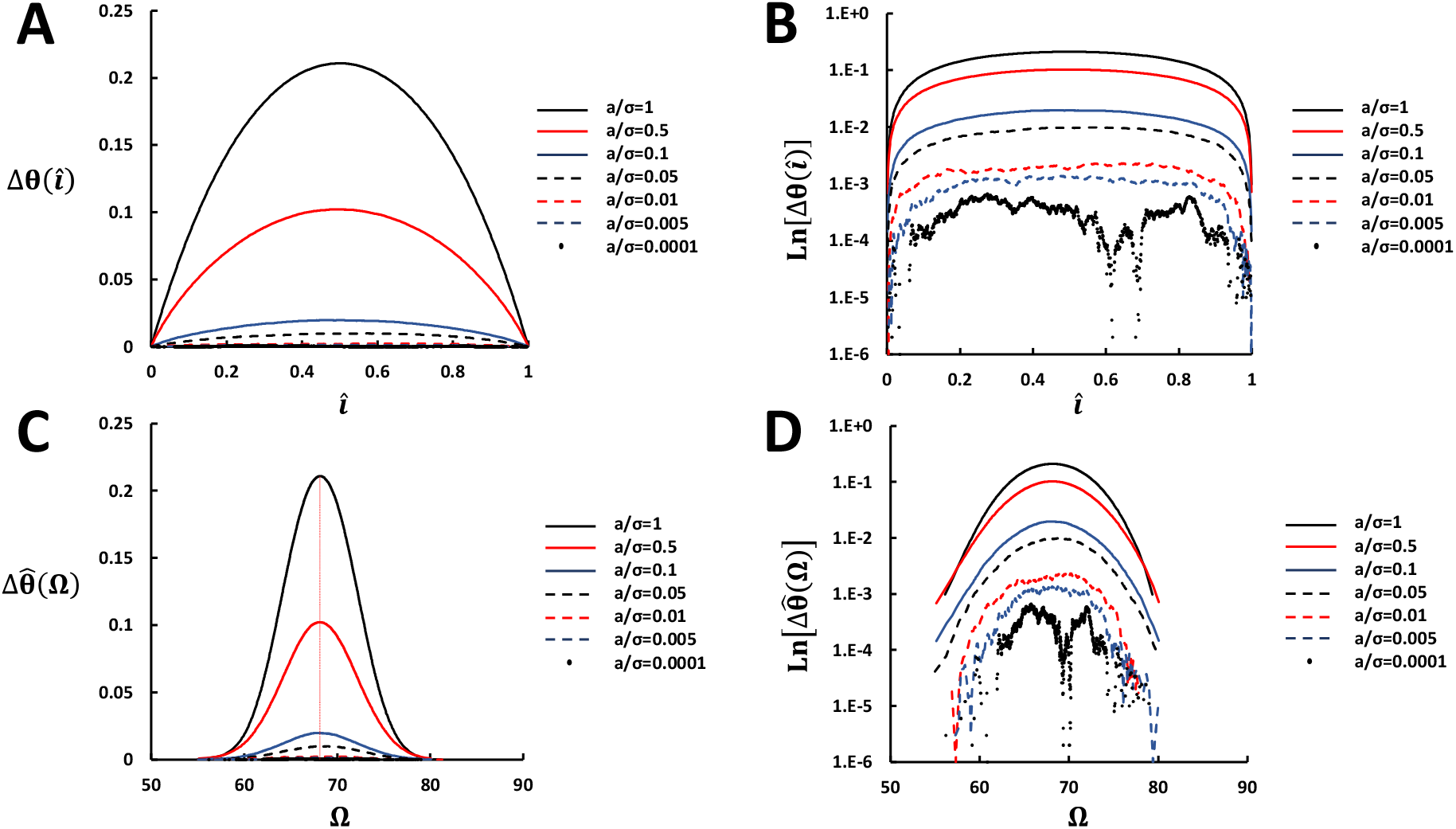
**(A)** Simulation of genetic paths difference in the space of positions as a function of log-scale values of the gene effect normalised by the phenotypic standard deviation, noted ‘*a*/*σ*’, for Δ*ω*_0_ = 0. **(B)** Natural-logarithm transformation of the genetic paths difference in the space of phenotype values as a function of log-scale values of the gene effect normalised by the phenotypic standard deviation, noted ‘*a*/*σ*’, for Δ*ω*_0_ = 0. **(C)** Simulations of genetic paths difference in the space of phenotype values as a function of log-scale values of the gene effect normalised by the phenotypic standard deviation, noted ‘*a*/*σ*’, for Δ*ω*_0_ = 0. **(D))** Natural-logarithm transformation of the genetic paths difference in the space of phenotype values as a function of log-scale values of the gene effect normalised by the phenotypic standard deviation, noted ‘*a*/*σ*’, for Δ*ω*_0_ = 0.

As shown in Figure 4B, the profile of the phenotype distribution density function is recovered with an amplitude that decreases as a/σ decreases. The red vertical dashed line in Figure 4D represents the mean phenotypic value. Using the natural logarithm to transform 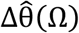 (Figure 4C) to 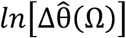 (Figure 4D) demonstrates that a difference between genetic paths can be seen for small gene effects.

One may then compare how perceptible the associations are using the new method by comparing Figures 1B and 1C (method of averages) and Figures 4D for identical allele frequencies and similar gene effects. Recall that GWAS rely on determining difference in averages (see Figure 1B or Figure 1C). However, the determination of a difference in the microstate averages rely on a strong gene effect (Figure 1B) or a very large population (see Figure 3) as otherwise the density functions of microstates collapse onto one. This is particularly visible when one compares the right-hand and left-hand graphs in figure 1B or Figure 1C. Thus, the results provided by Figure 4 suggest that GIFT can be applied to 1000 individuals to return information regarding potential genotype-phenotype associations that would not otherwise be possible, or extremely difficult, with current association studies. Concentrating on different allele frequencies given by p = 0.8 (Δω_0_ = 0.6) as an example. Figures 5A, and 5B are representations of 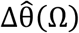 for log-scale values of a/σ using similar transformations as done for Δω_0_ = 0.

**Figure 5:**
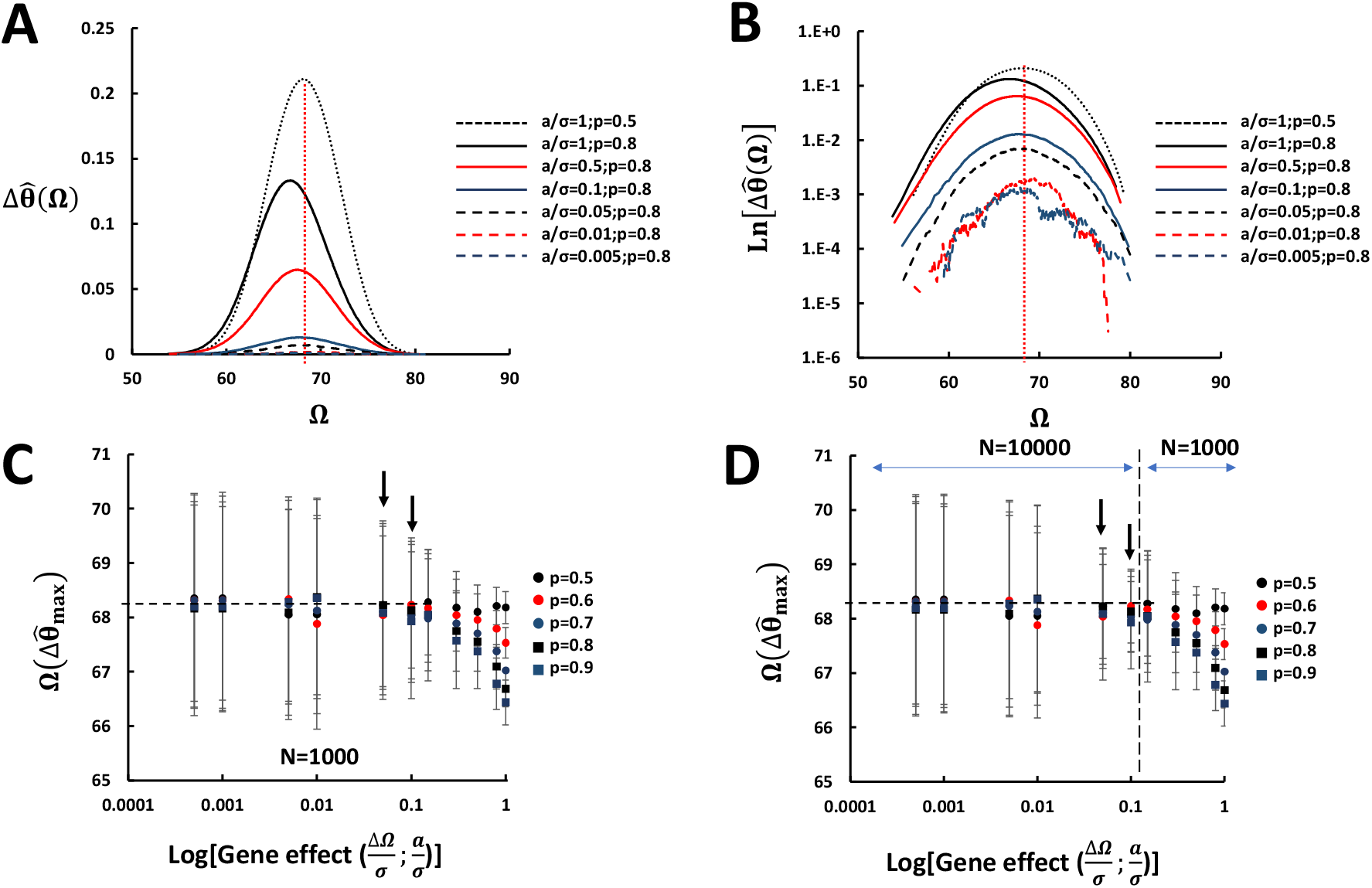
**(A)** Simulations of genetic paths difference in the space of phenotype values as a function of log-scale values of the gene effect normalised by the phenotypic standard deviation, noted ‘*a*/*σ*’, for Δ*ω*_0_ = 0.6. **(B)** Natural-logarithm transformation of the genetic paths difference in the space of phenotype values as a function of log-scale values of the gene effect normalised by the phenotypic standard deviation, noted ‘*a*/*σ*’, for Δ*ω*_0_ = 0.6. **(C)** Representation of the shift in the phenotype value for which the genetic path difference is maximal as a function of the gene effect ‘*a*/*σ*’ for different values of allele frequencies. N=1000 represents the number of individuals in the simulation. **(D)** Representation of the shift in the phenotype value for which the genetic paths difference is maximal when the population is increased by a factor 10 (N=10,000) for the gene effects verifying *a*/*σ* < 0.1.

Differences are clearly visible between Δω_0_ = 0 and Δω_0_ = 0.6, since the phenotype values for which 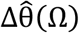’s are maximal have been shifted from the average value of the phenotype (indicated by the vertical red dashed line). This is not surprising because the simulation only imposed a set of genetic variances, without any constraint on the conservation of the average phenotype value.

However, the shift of the phenotype value for which 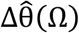 is maximal is of interest. As Eq.17 demonstrates that for small gene effects, the genetic path difference should be proportional to the phenotype distribution, that is, the phenotype value for which 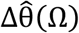 is extreme should be the average value of the phenotype.

Thus, to obtain a better visualisation of the impact of the gene effect on the positioning of the phenotype value 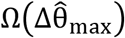 for which the genetic path difference is maximal, a set of simulations were also performed based on allele frequencies defined by p ∈ {0.5; 0.4; 0.3; 0.2; 0.1} for log-ranging values of gene effects (Figure 5C). Note that p ∈ {0.5; 0.6; 0.7; 0.8; 0.9} can be deduced from the symmetry around the average value of the phenotype.

Whilst the standard deviations obtained were not always negligible concerning 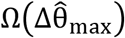, typically between 0.5 and 1 phenotypic standard deviation for small gene effects; Figure 5A demonstrated trends toward the average value of the phenotype with small gene effects. Indeed, below the simulated gene effect of a/σ~10^−l^, the average value of 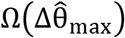 was remarkably similar to that of the average value of the phenotype, marked by the horizontal black dashed line.

To confirm this trend for small gene effects, that is, a/σ ≤ 0.1, we varied the population size from N = 10^3^ to N = 10^4^ to determine the presence of potential variations in 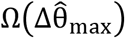 linked to the simulations. Results summarised in Figure 5B demonstrate that the only difference was a reduction in the standard deviations obtained for 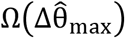 for the simulated gene effects comprised between 0.01 and 0.1 (see arrows Figures 5C and 5D pointing to different magnitude of the standard deviations). Namely, the initial symmetry of the phenotype distribution density function reappears, as expected (Eq.17).

Finally, Eq.17 suggests that for small gene effects 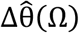 is proportional to the gene effect a/σ in the form, 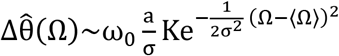. As a consequence it was decided to fit the all the curves 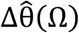 in Figures 4D and 5B with quadratic equations of the form 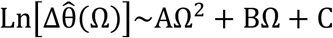; see Tables 2 and 3. Then, as 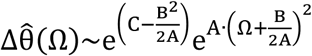 with such a fit, we expect by identification of Eq.17 that for small gene effects: 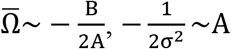 and 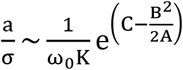. Tables 2 and 3 provide the estimations for both 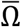 and σ and setting 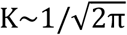, Figure 6 provides a comparison between the gene effect from the simulations, 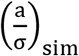, and the gene effect deduced from Eq.17, 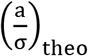.

**Table 2:** Nonlinear fit results for Figure 4C using 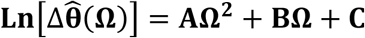.

|  | **a/$\sigma$=1** | **a/$\sigma$=0.5** | **a/$\sigma$=0.05** | **a/$\sigma$ =0.005** |
| --- | --- | --- | --- | --- |
|  | **p=0.5** | **p=0.5** | **p=0.5** | **p=0.5** |
| **A** | -0.0371 | -0.0321 | -0.0311 | -0.0435 |
| **B** | 5.0539 | 4.3799 | 4.2425 | 5.9241 |
| **C** | -173.69 | -151.48 | -149.5 | -208.35 |
| **r<sup>2</sup>** | 0.9960 | 0.9990 | 0.9937 | 0.8329 |
| **$\bar{\Omega}$** | **68.11** | **68.22** | **68.21** | **68.09** |
| **$\sigma^2$** | **13.48** | **15.58** | **16.08** | **11.49** |

**Table 3:** Nonlinear fit results for Figure 4F using 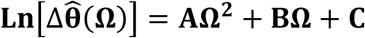.

|  | **a/$\sigma$=1**<br>**p=0.2** | **a/$\sigma$=0.5**<br>**p=0.8** | **a/$\sigma$=0.05**<br>**p=0.2** | **a/$\sigma$=0.005**<br>**p=0.2** |
| --- | --- | --- | --- | --- |
| **A** | -0.0318 | -0.0292 | -0.0282 | -0.0300 |
| **B** | 4.2666 | 3.9583 | 3.8514 | 4.0741 |
| **C** | -145.14 | -136.71 | -136.45 | -145.40 |
| **r<sup>2</sup>** | 0.9135 | 0.8892 | 0.8004 | 0.7554 |
| **$\bar{\Omega}$** | **67.08** | **67.77** | **68.28** | **67.90** |
| **$\sigma^2$** | **15.72** | **17.12** | **17.73** | **16.67** |

**Figure 6:**
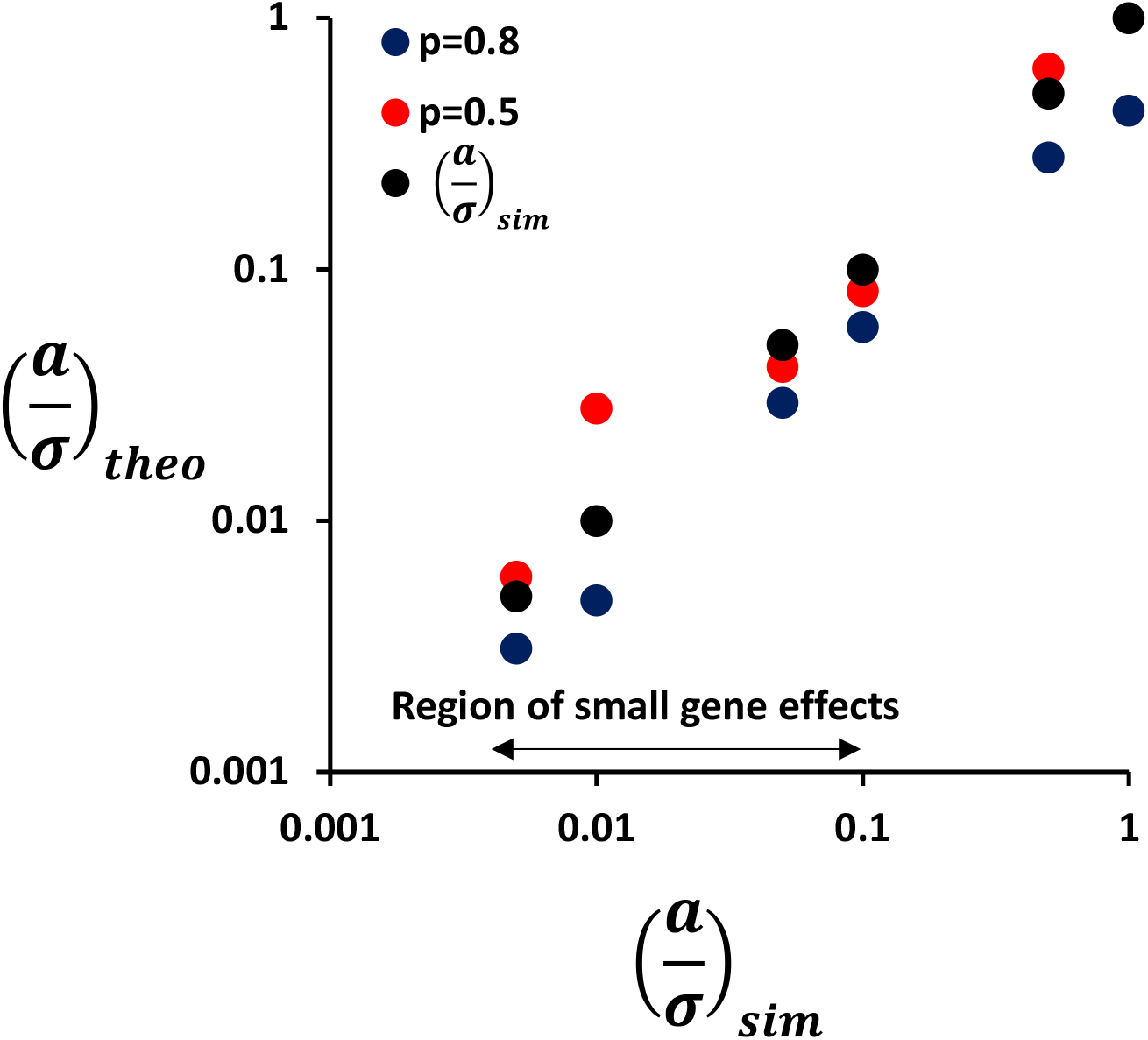
Comparison between simulated and theoretical gene effects the black dots correspond to the cases where the gene effects simulated are identical to the theoretical gene effects.

Thus, recalling that the phenotype average and variance of the population modelled are, respectively, 68.17 inch and 16.24 inch^2^; Table 2 and Table 3 demonstrate that fitting the genetic paths difference as a function of phenotype values with a quadratic curve recovers the magnitude of the average and variance of the phenotype used for the simulations for most log-scale values of the gene effect. Furthermore, the amplitude of 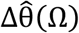 is also indicative of the gene effects involved.

## 5. Discussion

In his seminal paper (4), Fisher provided a synthesis between the genetic inheritance of continuous traits and the Mendelian scheme of inheritance using statistics and probability. His theory has become a landmark in genetics and heredity and its conceptual framework is still used today. While statistics is a natural field to employ when dealing with large datasets, the interpretation of data as well as the inferences that can be drawn from it rely fundamentally on probability density functions. As the act of creating categories to work with density function is acknowledging a sort of imprecision, our aim was to devise a different method ruling out the need for category. This new theoretical method, inspired by physics and named GIFT, uses the concept of phenotypic fields, and concentrate on ‘genetic paths’ to extract information on genotype phenotype mappings. It is then important to discuss the conceptual similarities and differences between GWAS and GIFT.

In term of conceptual similarities, we saw that the theory underscoring GIFT developed using Fisher’s assumptions recovers key concepts from quantitative genetics, including: (i) the Hardy-Weinberg coefficient locally, (ii) the Hardy-Weinberg coefficient at the population level, (iii) the gene effect, (iv) dominance, and (v) small gene effects involving common allele frequencies (30). In this context, GIFT and GWAS are similar. Finally, applying GIFT to simulated data based on Fisher’s assumption proved its sensitivity for extracting information on genotype-phenotype associations when sample sizes and gene effects were small. The reason for not considering the dominance in the simulations is linked to the fact that realistic genome-wide association studies have shown that with small effect sizes/small gene effect (which is the main area of concern of the current paper), dominance effects are often too small, and an additive model as suggested by Fisher works well enough (31).

In term of conceptual differences, three essential points can be discussed.

First of all, GIFT is more general that GWAS in the sense that the phenotypic fields can be any, namely do not have to be linear. The prescription of linear phenotypic fields in Fisher’s context comes from the symmetry associated with using the normal distribution as a template for any distribution density function, together with the assumption that the phenotypic and microstate variances are identical (4). When the constraint on the variances is released, the phenotypic fields become quadratic involving the variances as well as the averages. In this context GIFT has enabled us to define new parameters linked to microstate variances, that are, the pseudo-gene effect and the pseudo-dominance, which will probably help resolve controversies (20).

Secondly, in term of genetics what has been achieved so far is rather at odds with traditional ways of thinking about the notion of gene. Indeed, by defining the difference in genetic paths, Δ*θ*(Ω), one can say that it is the phenotype, i.e., phenotypic fields or information, that organises the configuration of genotypes and not the converse. In genetics the tradition is to think of genes as causing phenotypes. Here, a different way of thinking is suggested since it is the variation in phenotype values, resulting in our ability to generate a ranking process, that interacts with the microstates. Therefore, the phenotype is able to ‘select’ a set of genetic microstates. Recall that microstates ‘respond’ to, or interact with, the phenotypic fields only if they are associated with the phenotype. Consequently, this model suggests considering a genotype-phenotype ‘loop’, a.k.a. self-consistency. That is to say that if genes cause phenotypes (traditional view) and that phenotypes select gene microstates (present view), then an equivalence exists between phenotype and genotype. Supplementary Materials contain more information and development concerning the convergence and self-consistency of GIFT (SM6).

Finally, Fisher’s theory / GWAS has been built on considering the normal distribution. In general, ‘real’ density functions never come as normally distributed. Given that Fisher’s theory gives biological meanings to average and variance only, to define the ‘gene effect’ and ‘genetic/phenotypic variance’ linked to heredity, respectively; there is no biological meaning to any other statistical/mathematical parameters describing real density functions, such as for example the ‘skewness’. As GIFT uses curves, namely does not use average and variance as central parameters, this issue does not exist with GIFT. Said differently, GIFT frees GWAS from any preconceived idea of what statistics and probability applied to biology should be.

Taken as a whole, the work presented here is a first step suggesting that GIFT can be considered as a potential method for genotype-phenotype mappings. Supplementary Material SM7 contains more information concerning the signal-to-noise ratio when GIFT is used and SM8 (see Supplementary Materials) provides an initial illustration of the application of GIFT using real data based on GWAS results.

However the authors agree with the fact that more work needs to be done to compare GIFT to the vast literature concerning GWASes. For example, at present the model is quite simplistic in the way that, by construction, it does not allow the easy incorporation of covariates. Future works will relate covariates, such as age or sex, and for the case of human populations, genetic principal components (to account for population structure). In addition, GIFT will be compared against well-known statistical tests as used in GWAS (e.g., z-test/chi-square as parametric test, or the Kolmogorov–Smirnov test as non-parametric).

## 6. Conclusion

A century ago, Fisher devised a statistical method to map genotypes and phenotypes, which was essentially based on the measure of uncertainty. We present here a method taking as a paradigm the fact that certainty can exist with the possibility to measure phenotype and genotype with very high precision. In an associated paper, we present a theoretical methodology based on Shannon’s information enabling the significance of correlation using real genotype-phenotype data to be quantified (32). To conclude, this new method (GIFT) opens a new way to analyse genotype-phenotype mapping.

## Data accessibility

Code for the simulations presented in the manuscript can be found in the Supplementary Materials (see SM5). Genomic and phenotypic data analysed as part of this study in Supplementary Materials (see SM8) is already published and freely available online. The genomic data is part of the well-known Arabidopsis thaliana ‘1001 genomes’ project (https://doi.org/10.1186/gb-2009-10-5-107) and can be found online here: https://1001genomes.org/data-center.html. The Phenotype data was published in The Plant Journal (DOI: 10.1111/tpj.15177) and is freely available online via Ion Explorer here: https://bitbucket.org/ADAC_UoN/dr000081-web-service-ionome-seed-and-leaf-map/. The code used for the real-data analysis in SM8 in the Supplementary Materials can be found on Dryad, DOI: https://doi.org/10.5061/dryad.vx0k6djtp.

## Authors’ contributions

Conceptualisation of GIFT as a new method based on physics field theory: C.R.; theoretical developments: C.R., J.W.; simulation of genotype and phenotype data based on Fisher’s theory: C.R., Sa.Bl.; analysis of simulated or real data using GIFT: C.R., S.B., P.K.

## Competing interests

We have no competing interests.

## Funding

This work was supported by the Physics of Life Networks and The University of Nottingham.

## Acknowledgements

We thank Professors Andras Paldi, James Leigh, Patricia Harris, and Darren Logan for fruitful discussions concerning the urgent need to improve genotype-phenotype association methods for small sample sizes.

## Appendices

## Appendix 1: Differential expression of the genetic path in the space of phenotypic values

We assume that the phenotype values are measured precisely enough such that each individual has a unique phenotype value noted Ω_i_. As the population is composed of ‘N’ individuals one defines, 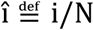 and Ω_î_, as the new position and its corresponding phenotype value, respectively. Thus Ω_l/N_ and Ω_1_ are the smallest and largest phenotype values, respectively.

As a result, the cumulative sum of presence probability of genetic microstates as a function of ‘î’ can be written as: 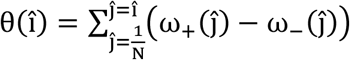, where 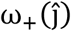 and 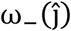 are the presence probabilities of microstates ‘+1’ and ‘−1’ at the position 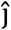. Using the continuum limit, one deduces also, 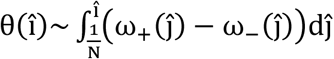.

As the ranking of phenotype values was introduced to define, θ(î), the genetic path can also be expressed as a function of phenotype values under the form: 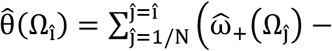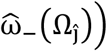 where the hat 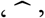 is added to describe new functions linked to phenotypic values. Accordingly, by definition, 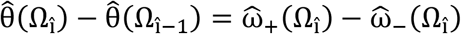. Assuming that the measured phenotype values are sufficiently close, the continuum limit can be used to determine 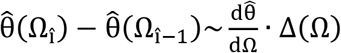, where Δ(Ω) is the spacing between two consecutive individuals in the space of phenotype values. One deduces then: 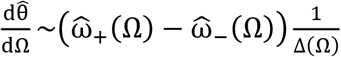.

As a result, the genetic path can be expressed in the space of the phenotype values as:

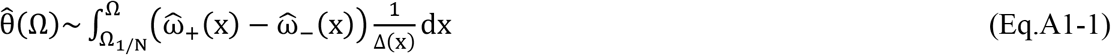

Where 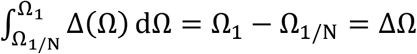. As Δ(Ω) is the phenotypic space between individual, provided that the population is large and dense enough one can relate ‘Δ(Ω)’ to the phenotype distribution density function, P_Ω_(Ω), under the form 1/Δ(Ω)~P_Ω_(Ω) (SM1 in the Supplementary Materials). The different elements that lead to the formation of a genetic path can now be addressed.

## Appendix 2: Entropy of the string of microstates

Keeping the notations 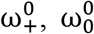 and 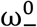, for the genetic microstate frequencies of a given genome position across the population of individuals, we aim to determine the expressions of ω_+_(î), ω_0_(î) and ω_−_(î) given the information obtained upon ordering the genotypes as a function of phenotype values along the î-axis.

The default genetic path, θ_0_, as a straight line is defined by the absence of information on phenotype values, which is similar to an absence of association between the genetic microstates and the phenotype values, leading to an apparent disordering of genetic microstates. One way to measure this disordering is by using the ‘entropy’ of the string of genetic microstates for the genome position considered. In this context, the entropy is given by calculating the number of possible combinations of placing 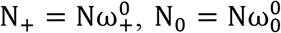 and 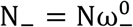 genetic microstates over ‘N’ possible positions. Consequently, the entropy of the default genetic path is S_0_ = N!/N_+_!N_0_!N_−_!; and for ‘N’ large enough using Stirling’s formula one deduces: 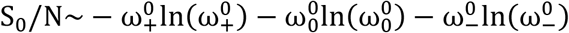, that can be rewritten in the continuum limit as: 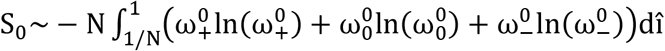. Note that, as the genetic microstate frequencies are constant in this case, the entropy can be rewritten using the phenotype values as,

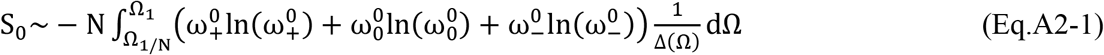

When information about phenotype values and their ranking is given and when the genome position considered is associated with the phenotype, S_0_ is transformed to S where:

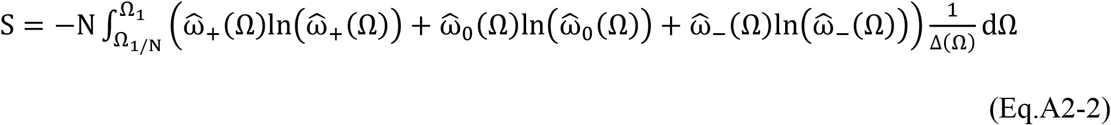

As a result, the entropy difference, S − S_0_ when non-null provides information on whether the genome position is associated with phenotype values. Thus the difference, S − S_0_, can be thought of as a ‘transformation’ in a physical/thermodynamic sense. That is, the difference in entropies must be balanced by a term that is linked to the association (or interaction) between the genetic microstates and the phenotype values.

## Appendix 3: Interaction energy between microstates and subfields

As the difference, S − S_0_, is linked to the information gained from knowing phenotypic values and ranking them (Appendix 2), given the existence of three distinct genetic microstates, one can define three distinct functions a.k.a. phenotypic fields ‘u_+_(Ω)’, ‘u_0_(Ω)’ and ‘u_−_(Ω)’ that are fundamentally related to changes in the phenotype-associated genetic path. In this context, the entire genetic path can be defined with a function representing the sum of interactions between each of the genetic microstates and phenotypic fields under the form:

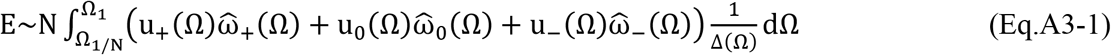

In this context, one may consider that the set of microstates changes the configuration because the fields are ‘switch on’. This implies that for the genome positions that are not involved in the formation of the phenotype considered, the switch does not work, that is, the fields are null. In this context, one can consider the equivalence, S − S_0_~E. As a result, the relationship to optimise is: ΔS − E = 0.

## Appendix 4: Optimisation of ΔS − E

Recalling the conservation of genetic microstates for the genome position considered: 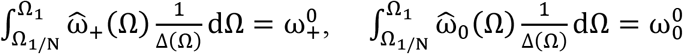 and 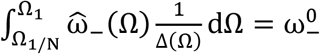 together with the conservation of probability, 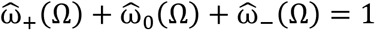, Euler-Lagrange’s method can then be used to determine the optimal configuration for 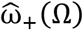, 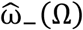 and 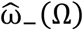 in a context where the phenotypic fields are imposed. By defining α_+_, α_0_ and α_−_, the Lagrange multipliers for the conservation of genetic microstates, the relation to optimise with regard to the genetic microstate frequencies 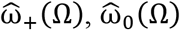 and 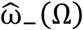 is then,

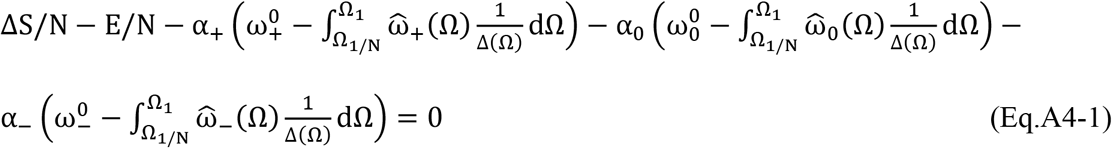

Using the conservation of genetic microstate frequencies, 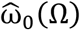, can be replaced by, 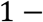 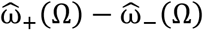, and a variational calculus can be performed on the genetic microstate frequencies, leading to two conditions:

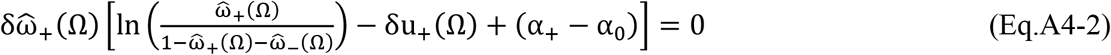

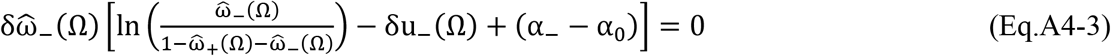

Where 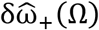 and 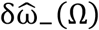 are small variations in the presence probabilities of microstates ‘+1’ and ‘−1’, and 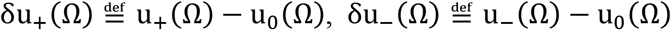. Finally, 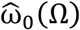 can be deduced using 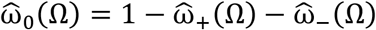. Using the conditions 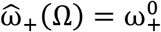, 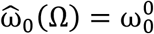 and 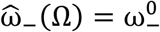 when the fields are null, one obtains,

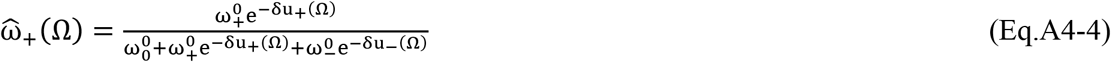

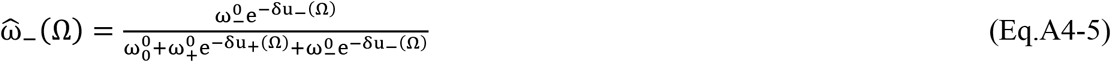

To make the asymmetries of the problem more apparent, the following are defined for genetic microstates: 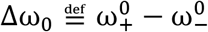 and 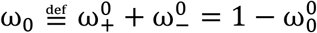; and for the phenotypic fields: 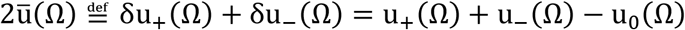 and 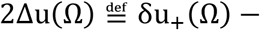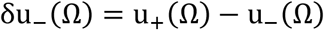. Then, the difference and sum of 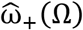 and 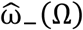 can be rewritten as follow:

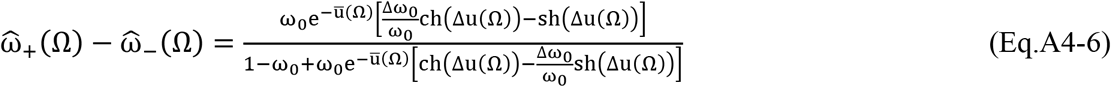

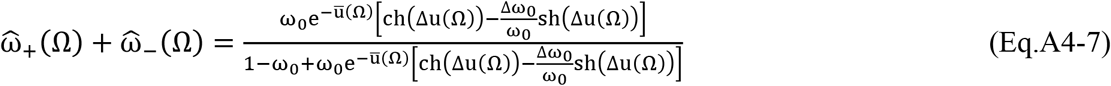

Noting that: −1 ≤ Δω_0_/ω_0_ ≤ +1, a new phenotype value is defined and noted ‘Ω_0_’ by setting 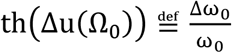. Then, the difference and sum of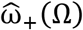 and 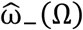 can be rewritten as:

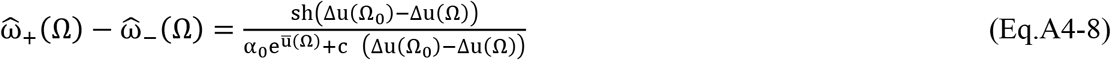

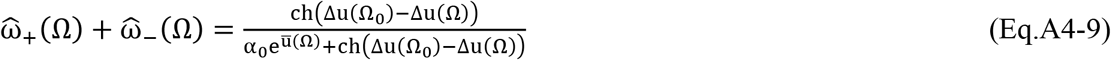

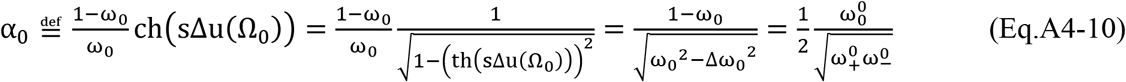

The new variable ‘Ω_0_’ is the phenotype value corresponding to the condition 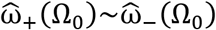. The meaning of the constant ‘α_0_’ can be related to the Hardy-Weinberg law. Hardy-Weinberg law based on random mating in a population provides a relationship between the genetic microstate frequencies under the form: p^2^ + 2pq + q^2^ = 1, where p^2^ and q^2^ are the genotype frequencies of genetic microstates ‘+1’ and ‘−1’, i.e. homozygote genotypes aa and AA, respectively; and 2pq the genotype frequency for genetic microstate ‘0’, i.e. the heterozygote genotype Aa. In our case, this corresponds to replacing p^2^, q^2^ and 2pq with, respectively, 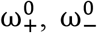 and 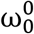. Consequently, the Hardy-Weinberg law imposes α_0_ = 1 with α_0_ ≠ 1 corresponding to a deviation from the law. However, this term is expected to remain stable upon any changes of allele or genotype frequencies suggesting therefore that, genetically, any changes in ‘Δω_0_’ are to some extent compensated by corresponding changes in ‘ω_0_’.

We can now turn to the full expression of the genetic path difference in the space of the phenotype value:

## Appendix 5: Expression of the difference between the phenotype responding and default genetic paths expressed in the phenotypic space and conservation of genetic microstate frequencies

The phenotype-associated genetic path is simply the integration of (Eq.A4-8) over the phenotype values that is given, as seen above (Eq.A1-1 in Appendix 1), by: 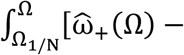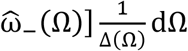. The default genetic path is deduced from considering that the difference in the presence probabilities between the genetic microstates ‘+1’ and ‘−1’ is constant, i.e.: 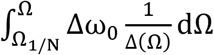. By rewriting ‘Δω_0_’ as ‘ω_0_(Δω_0_/ω_0_)’, where Δω_0_/ω_0_ = th(Δu(Ω_0_)) and deducing ‘ω_0_’ from α_0_ = (1 − ω_0_)ch(Δu(Ω_0_))/ω_0_; it follows: 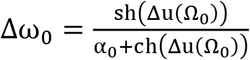. As a result, the difference between the phenotype-associated and default genetic paths expressed as a function of phenotypic fields within the continuum limit is:

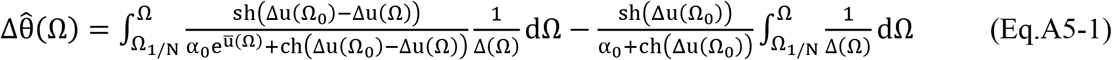

The conservation of genetic microstates needs to be added regardless of the genetic path taken, that is, 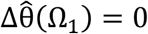, expressed as:

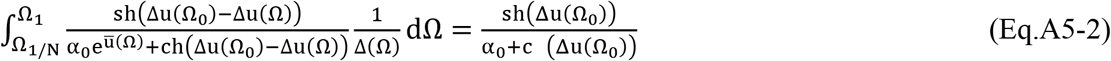

## Notes

### Competing Interest Statement

The authors have declared no competing interest.

### Summary of Updates

Few typos were spotted on by one of the referee (Minor changes prior to publication).

## References

1. Buniello A, MacArthur JAL, Cerezo M, Harris LW, Hayhurst J, Malangone C, et al. The NHGRI-EBI GWAS Catalog of published genome-wide association studies, targeted arrays and summary statistics 2019. Nucleic Acids Res. 2019 Jan;47(D1):D1005–12.

2. Sudlow C, Gallacher J, Allen N, Beral V, Burton P, Danesh J, et al. UK Biobank: An Open Access Resource for Identifying the Causes of a Wide Range of Complex Diseases of Middle and Old Age. PLOS Med [Internet]. 2015 Mar 31;12(3):e1001779. Available from: https://doi.org/10.1371/journal.pmed.1001779

3. Smith BH, Campbell A, Linksted P, Fitzpatrick B, Jackson C, Kerr SM, et al. Cohort Profile: Generation Scotland: Scottish Family Health Study (GS:SFHS). The study, its participants and their potential for genetic research on health and illness. Int J Epidemiol. 2013 Jun;42(3):689–700.

4. Fisher RA. XV.—The Correlation between Relatives on the Supposition of Mendelian Inheritance. Trans R Soc Edinburgh. 1919;52(2):399–433.

5. Fisher RA. XXI.—On the Dominance Ratio. Proc R Soc Edinburgh [Internet]. 2014/09/15. 1923;42:321–41. Available from: https://www.cambridge.org/core/article/xxion-the-dominance-ratio/2CDBE1D374EBE9B2774DF301BA98A584

6. Visscher PM, Goddard ME. From R.A. Fisher’s 1918 Paper to GWAS a Century Later. Genetics. 2019 Apr;211(4):1125–30.

7. Hivert V, Wray NR, Visscher PM. Gene action, genetic variation, and GWAS: A user-friendly web tool. PLoS Genet. 2021 May;17(5):e1009548.

8. Stephens M, Balding DJ. Bayesian statistical methods for genetic association studies. Nat Rev Genet [Internet]. 2009;10(10):681–90. Available from: https://doi.org/10.1038/nrg2615

9. Beaumont MA, Rannala B. The Bayesian revolution in genetics. Nat Rev Genet. 2004 Apr;5(4):251–61.

10. Boyle EA, Li YI, Pritchard JK. An Expanded View of Complex Traits: From Polygenic to Omnigenic. Cell [Internet]. 2017;169(7):1177–86. Available from: https://www.sciencedirect.com/science/article/pii/S0092867417306293

11. van der Harst P, Verweij N. Identification of 64 Novel Genetic Loci Provides an Expanded View on the Genetic Architecture of Coronary Artery Disease. Circ Res. 2018 Feb;122(3):433–43.

12. Herrmann M, Yampolsky LY. False and true positives in arthropod thermal adaptation candidate gene lists. Genetica. 2021 Jun;149(3):143–53.

13. Mähler N, Schiffthaler B, Robinson KM, Terebieniec BK, Vučak M, Mannapperuma C, et al. Leaf shape in Populus tremula is a complex, omnigenic trait. Ecol Evol. 2020 Nov;10(21):11922–40.

14. Zhang W, Reeves GR, Tautz D. Testing Implications of the Omnigenic Model for the Genetic Analysis of Loci Identified through Genome-wide Association. Curr Biol. 2021 Mar;31(5):1092-1098.e6.

15. Vuckovic D, Bao EL, Akbari P, Lareau CA, Mousas A, Jiang T, et al. The Polygenic and Monogenic Basis of Blood Traits and Diseases. Cell. 2020 Sep;182(5):1214-1231.e11.

16. Mathieson I. The omnigenic model and polygenic prediction of complex traits. Am J Hum Genet. 2021 Sep;108(9):1558–63.

17. Galton F. Regression Towards Mediocrity in Hereditary Stature. J Anthropol Inst Gt Britain Irel [Internet]. 1886 Oct 22;15:246–63. Available from: http://www.jstor.org/stable/2841583

18. Visscher PM, Medland SE, Ferreira MAR, Morley KI, Zhu G, Cornes BK, et al. Assumption-free estimation of heritability from genome-wide identity-by-descent sharing between full siblings. PLoS Genet. 2006 Mar;2(3):e41.

19. Silventoinen K, Sammalisto S, Perola M, Boomsma DI, Cornes BK, Davis C, et al. Heritability of adult body height: a comparative study of twin cohorts in eight countries. Twin Res. 2003 Oct;6(5):399–408.

20. Nelson RM, Pettersson ME, Carlborg Ö. A century after Fisher: time for a new paradigm in quantitative genetics. Trends Genet. 2013 Dec;29(12):669–76.

21. Yang J, Benyamin B, McEvoy BP, Gordon S, Henders AK, Nyholt DR, et al. Common SNPs explain a large proportion of the heritability for human height. Nat Genet [Internet]. 2010/06/20. 2010 Jul;42(7):565–9. Available from: https://pubmed.ncbi.nlm.nih.gov/20562875

22. Wood AR, Esko T, Yang J, Vedantam S, Pers TH, Gustafsson S, et al. Defining the role of common variation in the genomic and biological architecture of adult human height. Nat Genet. 2014 Nov;46(11):1173–86.

23. Visscher PM, Hill WG, Wray NR. Heritability in the genomics era–-concepts and misconceptions. Nat Rev Genet. 2008 Apr;9(4):255–66.

24. Yengo L, Vedantam S, Marouli E, Sidorenko J, Bartell E, Sakaue S, et al. A Saturated Map of Common Genetic Variants Associated with Human Height from 5.4 Million Individuals of Diverse Ancestries. bioRxiv [Internet]. 2022; Available from: https://www.biorxiv.org/content/early/2022/01/10/2022.01.07.475305

25. Stigler SM. The History of Statistics: The Measurement of Uncertainty before 1900 [Internet]. Harvard University Press; 1990. Available from: https://books.google.co.uk/books?id=nWn9DwAAQBAJ

26. Macdonald A, Hawkes LA, Corrigan DK. Recent advances in biomedical, biosensor and clinical measurement devices for use in humans and the potential application of these technologies for the study of physiology and disease in wild animals. Philos Trans R Soc London Ser B, Biol Sci. 2021 Aug;376(1831):20200228.

27. Falconer DS. Introduction to quantitative genetics. Harlow, England: Prentice Hall; 1996.

28. Lonsdale J, Thomas J, Salvatore M, Phillips R, Lo E, Shad S, et al. The Genotype-Tissue Expression (GTEx) project. Nat Genet [Internet]. 2013;45(6):580–5. Available from: https://doi.org/10.1038/ng.2653

29. Delongchamp R, Faramawi MF, Feingold E, Chung D, Abouelenein S. The Association between SNPs and a Quantitative Trait: Power Calculation. Eur J Environ public Heal. 2018;2(2).

30. Park J-H, Gail MH, Weinberg CR, Carroll RJ, Chung CC, Wang Z, et al. Distribution of allele frequencies and effect sizes and their interrelationships for common genetic susceptibility variants. Proc Natl Acad Sci [Internet]. 2011;108(44):18026–31. Available from: https://www.pnas.org/content/108/44/18026

31. Sham PC, Cherny SS, Purcell S, Hewitt JK. Power of Linkage versus Association Analysis of Quantitative Traits, by Use of Variance-Components Models, for Sibship Data. Am J Hum Genet [Internet]. 2000;66(5):1616–30. Available from: https://www.sciencedirect.com/science/article/pii/S0002929707629929

32. Wattis J, Bray S, Kyratzi P, Rauch C. Analysis of Genotype-Phenotype Association using Fields and Information Theory. arXiv Prepr arXiv220211989. 2022;

