## Supplementary Materials for "GIFT: New method for the genetic analysis of small gene effects involving small sample sizes" (from another version of this preprint: version 3, 2022-09-15)

**Key words: phenotype-genotype mapping, fields theory, complex traits, GWAS, GIFT**

**CONTENTS:**

**Page2 - SM1: Determination of the relationship between '$\Delta(\Omega)$' and ' $P_{\Omega}(\Omega)$'.**

**Page 4 - SM2: Minimalist model for phenotypic fields.**

**Page 7 - SM3: Fisher fields.**

**Page 8 - SM4: Condition imposed by the conservation of genetic microstate frequencies when the minimalist model (Fisher fields) is considered.**

**Page 15 - SM5: Code for the simulations**

**Page 21 - SM6: Convergence and Self-consistency of GIFT.**

**Page 27 – SM7: Signal-to-noise ratio when GIFT is used.**

**Page 30 – SM8: GIFT using real data.**

**SM1: Determination of the relationship between '$\Delta(\Omega)$' and ' $P_\Omega(\Omega)$'**

To provide a description of '$\Delta(\Omega)$' in relation to the phenotype distribution density function '$NP_\Omega(\Omega)$' as usually used in genotype-phenotype association studies, we recall that '$NP_\Omega(\Omega)$' is defined by frequentist probabilities as is the number of individuals, '$\delta i$', having a phenotype value comprised between '$\Omega - \delta\Omega/2$' and '$\Omega + \delta\Omega/2$, i.e.: $NP_\Omega(\Omega) = \delta i/\delta\Omega$ .

As a result, '$1/NP_\Omega(\Omega) \sim \delta\Omega/\delta i$' can be assimilated into the free average phenotypic space available per individual present within a category.

Using '$\Delta(\Omega)$', the free average space per individual can also be written as $\langle\Delta(\Omega)\rangle \sim \frac{\delta\Omega}{\delta i} \sim \frac{1}{\delta\Omega}\int_{\Omega-\delta\Omega/2}^{\Omega+\delta\ /2} \Delta(x)\, dx$.

Likewise, the number of individuals with a phenotype value comprised between '$\Omega - \delta\Omega/2$' and '$\Omega + \delta\Omega/2$' can be expressed by, $\delta i = \int_{\Omega-\delta\Omega/2}^{\Omega+\delta\Omega/2} \frac{dx}{\Delta(x)}$, from which it is deduced $\langle\frac{1}{\Delta(\Omega)}\rangle \sim \frac{\delta i}{\delta\Omega} \sim \frac{1}{\delta\Omega}\int_{\Omega-\delta\Omega/2}^{\Omega+\delta\Omega/2} \frac{dx}{\Delta(x)}$. As a result, provided that '$\Delta(\Omega)$' is determined, information on the phenotype distribution density function can be inferred if needed.

A more extreme approach is also considered. Let us assume three consecutive individuals and that the first individual has a phenotype value '$\Omega_1$'. Then the second individual is expected to have a phenotype value $\Omega_2 = \Omega_1 + \Delta(\Omega_1)$. Similarly, the third individual must have the phenotype value $\Omega_3 = \Omega_2 + \Delta(\Omega_2) = \Omega_1 + \Delta(\Omega_1) + \Delta(\Omega_1 + \Delta(\Omega_1))$. Let us assume the existence of a function $f_\Omega(\Omega)$, that determines the exact free phenotype space per individual as a function of the phenotype values. The free volume for the second individual can then be defined as: $f_\Omega(\Omega_2) \stackrel{\text{def}}{=} \frac{1}{2}(\Omega_3 - \Omega_2) + \frac{1}{2}(\Omega_2 - \Omega_1)$. As a result, by using the recurring relations between '$\Omega_1$', '$\Omega_2$' and '$\Omega_3$' it can be deduced that the free phenotype space per individual as a function of its phenotype value can be written as: $2f_\Omega(\Omega + \Delta(\Omega)) = \Delta(\Omega + \Delta(\Omega)) + \Delta(\Omega)$. The variable $1/f_\Omega(\Omega)$ is then similar to the concentration of a single individual in the space of the phenotype values. This relation can then be directly related to the definition, $NP_\Omega(\Omega) =$

$\delta i/\delta\Omega$, by dividing the position by the total number of individuals, resulting in $P_\Omega(\Omega) = \delta\hat{\imath}/\delta\Omega$, where '$P_\Omega(\Omega)$' is then also the concentration of a single individual in the space of phenotype value.

Thus, in the ideal case where a large population has been genotyped and phenotyped and that the difference between the phenotype values of two consecutive individuals is much smaller than their respective phenotype value, i.e. when $\Omega/\Delta(\Omega) \ll 1$; the concentration of a single individual in the space of phenotype values can be approached by: $\frac{1}{f_\Omega(\Omega)} \sim P_\Omega(\Omega) \sim \frac{1}{\Delta(\Omega)}$.

Finally, whilst '$\Delta(\Omega)$' is strictly defined within the interval $[\Omega_{1/N}, \Omega_1]$, this interval may be extended artificially by defining $1/\Delta(\Omega) \sim 0$ outside this interval. This assumption is equivalent to stating that the distance between individuals in the phenotype space is infinitely large for extreme phenotype values. Such an assumption is coherent with the asymptotic limits of '$P_\Omega(\Omega)$' in this case.

**SM2: Minimalist model for phenotypic fields**

As '$u_0(\Omega)$' is the reference field, the simplest model consists in considering the reference field set to zero and the fields '$u_+(\Omega)$' and '$u_-(\Omega)$' acting anti symmetrically on the microstate '+1' and '-1' such that: $u_+(\Omega) + u_-(\Omega) = 0$. This condition guarantees the migration of microstate '+1' and '-1' in opposite directions between the two configurations aforementioned (see Fig.2 main text).

Thus, setting, $u(\Omega) \stackrel{\text{def}}{=} \frac{1}{2}u_+(\Omega) = -\frac{1}{2}u_-(\Omega)$, Eq.8 (main text) is reduced to:

$$\int_{\Omega_{1/N}}^{\Omega_1} \frac{sh(u(\Omega_0)-u(\Omega))}{\alpha_0+ch(u(\Omega_0)-u(\Omega))}\frac{1}{\Delta(\Omega)}d\Omega = \frac{sh(u(\Omega_0))}{\alpha_0+ch(u(\Omega_0))}$$

To determine the expression of '$u(\Omega)$', one notes that by using Eq.3 and Eq.4 from the main text: $\widehat{\omega}_+(\Omega) \to 1$ (resp. $\widehat{\omega}_-(\Omega) \to 1$) if $u(\Omega) \to -\infty$ (resp. $u(\Omega) \to +\infty$), suggesting that the field needs to change sign, that is, is an odd function of the phenotype values. Let us note '$\Omega_m$' the phenotype value verifying $u(\Omega_m){\sim}0$. Without loss of generality, the field can be rewritten in the form $u(\Omega - \Omega_m)$ with $u(0){\sim}0$ and consequently, $u(\Omega_0) = u(\Omega_0 - \Omega_m)$. The derivative of Eq.7 from the main text is thus given by:

$$\frac{d\Delta\widehat{\theta}}{d\Omega} = \left[\frac{sh(u(\Omega_0-\Omega_m)-u(\Omega-\Omega_m))}{\alpha_0+ch(u(\Omega_0-\Omega_m)-u(\Omega-\Omega_m))} - \frac{sh(u(\Omega_0-\Omega_m))}{\alpha_0+c\ \ (u(\Omega_0-\Omega_m))}\right]\frac{1}{\Delta(\Omega)}$$

The latter expression demonstrates that '$\Omega_m$' is the phenotype value for which the genetic paths difference attains its extreme value irrespective of the gene microstate frequencies involved. To determine the signification of '$\Omega_m$' let us return in the space of positions (Eq.1 in main text) and rewrite the derivative of the phenotype-associated genetic path as:

$$\frac{d\theta}{d\hat{\imath}} = \frac{sh\left(u(\Omega_{\hat{\imath}_0}-\Omega_{\hat{\imath}_m})-u(\Omega_{\hat{\imath}}-\Omega_{\hat{\imath}_m})\right)}{\alpha_0+ch\left(u(\Omega_{\hat{\imath}_0}-\Omega_{\hat{\imath}_m})-u(\Omega_{\hat{\imath}}-\Omega_{\hat{\imath}_m})\right)}$$

One notes that the phenotype-associated genetic path is extreme, that is, $d\theta/d\hat{\imath}{\sim}0$, when the argument contained in the hyperbolic sine function is null. Three positions are defined in this argument, i.e., '$\hat{\imath}_0$', '$\hat{\imath}_m$' and '$\hat{\imath}$'. The signification of '$\Omega_{\hat{\imath}_m}$' can be discussed in the particular case where $\Delta\omega_0 = 0$, i.e., when the number of genetic microstates '+1' and '-1' are equal. In this case, as $\alpha_0 = \frac{1-\omega_0}{\omega_0}\mathrm{ch}\left(\Delta u\left(\Omega_{\hat{\imath}_0} - \Omega_{\hat{\imath}_m}\right)\right)$:

$$\Delta\omega_0 = \frac{\mathrm{sh}\left(u\left(\Omega_{\hat{\imath}_0} - \Omega_{\hat{\imath}_m}\right)\right)}{\alpha_0 + \mathrm{ch}\left(u\left(\Omega_{\hat{\imath}_0} - \Omega_{\hat{\imath}_m}\right)\right)} = \omega_0 \mathrm{th}\left(u\left(\Omega_{\hat{\imath}_0} - \Omega_{\hat{\imath}_m}\right)\right)$$

As a result, $\Delta\omega_0 = 0$ implies $u\left(\Omega_{\hat{\imath}_0} - \Omega_{\hat{\imath}_m}\right) \sim 0$. Therefore, due to the symmetry involved when $\Delta\omega_0 = 0$ and the fact that the field acts anti-symmetrically on the genetic microstates '+1' and '-1' the phenotype-associated genetic path would attain its extreme value at the median position. Thus, '$\Omega_{\hat{\imath}_m}$' or equivalently, '$\Omega_m$', corresponds to the phenotype value at the median position, '$\hat{\imath}_m$' (see Figure SM2-A below).

Now, consider the case $\Delta\omega_0 \neq 0$. In this case $u\left(\Omega_{\hat{\imath}_0} - \Omega_{\hat{\imath}_m}\right) \neq 0$, and the phenotype-associated genetic path would attain its extreme value at the position '$\hat{\imath}_0$'. However, as the field acts anti-symmetrically on the genetic microstates '+1' and '-1' it is always possible to isolate a subpopulation from the initial population involving a subset of ranked phenotype values such that the new phenotype-associated genetic path is, in the subpopulation, without asymmetry. Thus, in the subpopulation the new genetic path would be symmetrical and maximal for a phenotype value at the position '$\Omega_{\hat{\imath}'_m}$', corresponding to '$\Omega_{\hat{\imath}_0}$' in the initial population and where '$\hat{\imath}'_m$' would be a new median position. As a result, writing $u\left(\Omega_{\hat{\imath}} - \Omega_{\hat{\imath}_m}\right) - u\left(\Omega_{\hat{\imath}_0} - \Omega_{\hat{\imath}_m}\right) \sim 0$ for the entire population or $u\left(\Omega_{\hat{\imath}} - \Omega_{\hat{\imath}'_m}\right) = u\left(\Omega_{\hat{\imath}} - \Omega_{\hat{\imath}_0}\right) \sim 0$ for the selected subpopulation is equivalent (see Figure SM2-B). This implies a rule of additivity for the phenotypic field, expressed as a function of phenotype values, $u\left(\Omega_{\hat{\imath}} - \Omega_{\hat{\imath}_m}\right) - u\left(\Omega_{\hat{\imath}_0} - \Omega_{\hat{\imath}_m}\right) \sim u\left(\Omega_{\hat{\imath}} - \Omega_{\hat{\imath}_0}\right)$, leading to the conclusion that the field is a linear function of phenotype values.

As a result, the field is expressed as: $u(\Omega - \Omega_m) \sim s(\Omega - \Omega_m)$ where 's' is a constant and the genetic paths difference can be rewritten as:

$$\Delta\hat{\theta}(\Omega) \sim \int_{\Omega_{1/N}}^{\Omega} \left(\frac{\mathrm{sh}(s(\Omega_0 - \Omega))}{\alpha_0 + \mathrm{ch}(s(\Omega_0 - \Omega))} - \frac{\mathrm{sh}(s(\Omega_0 - \Omega_m))}{\alpha_0 + \mathrm{ch}(s(\Omega_0 - \Omega_m))}\right) \frac{1}{\Delta(\Omega)} d\Omega \qquad \text{(Eq.SM.2-1)}$$

This minimalist model implies that phenotypic fields must be linear.

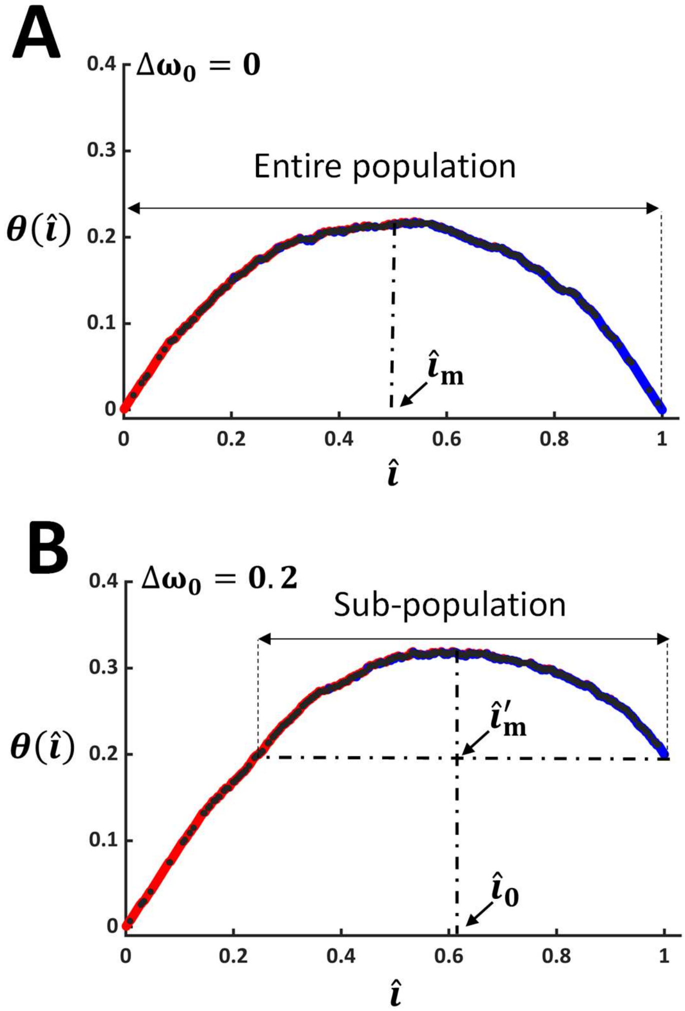


**Figure SM2:** Genetic paths modelled using GWAS when the dominance is null. The plots represent the cumulative sum of microstates for two different genome positions characterised by $\Delta\omega_0 = 0$ **(A)** and $\Delta\omega_0 = 0.2$ **(B)**. The microstates are colour coded as represented in Figures 1 and 2. Contrary to the symmetrical case given by $\Delta\omega_0 = 0$ and where the maximal value of the genetic path expressed in the space of positions occurs at $\hat{\imath}_m = 0.5$; when $\Delta\omega_0 = 0.2$ the asymmetry in the genetic microstates operates a translation of the phenotype field implying that the maximal value of the genetic path is shifted to around $\hat{\imath}_0 \sim 0.6$. Within the framework of GIFT this corresponds to the definition, in the space of positions, of the phenotype value '$\Omega_0$'. It is interesting to note that using simulations a subpopulation can be defined, namely that similarities exist between different genetic microstate frequencies pertaining to the fact that the fields for '+1' and '-1' are symmetrical when no dominance is involved (see Equations 12).

**SM3: Fisher fields**

The identity between the two versions of GIFT, that is, coarse-grained or not, can be established by noting that for each microstate, the number for the category concerned can also be expressed as:

$$\delta N_+ = \int_{\Omega}^{\Omega+\delta\Omega} \hat{\omega}_+(\Omega)\frac{1}{\Delta(\Omega)}d\Omega \sim \hat{\omega}_+(\Omega)\frac{\delta\Omega}{\Delta(\Omega)} + O(\delta\Omega^2)$$

$$\delta N_0 = \int_{\Omega}^{\Omega+\delta\Omega} \hat{\omega}_0(\Omega)\frac{1}{\Delta(\Omega)}d\Omega \sim \hat{\omega}_0(\Omega)\frac{\delta\Omega}{\Delta(\Omega)} + O(\delta\Omega^2)$$

$$\delta N_- = \int_{\Omega}^{\Omega+\delta\Omega} \hat{\omega}_-(\Omega)\frac{1}{\Delta(\Omega)}d\Omega \sim \hat{\omega}_-(\Omega)\frac{\delta\Omega}{\Delta(\Omega)} + O(\delta\Omega^2)$$

As the ratio '$\delta\Omega/\Delta(\Omega)$' corresponds to the number of individuals within the category, that is, $\delta\Omega/\Delta(\Omega) \sim \delta N \sim NP_\Omega(\Omega)\delta\Omega$, by using the distribution density functions for each microstate, that is, $\delta N_+ \sim N_+^0 \cdot P_+(\Omega)\delta\Omega$, $\delta N_0 \sim N_0^0 \cdot P_0(\Omega)\delta\Omega$ and $\delta N_- \sim N_-^0 \cdot P_-(\Omega)\delta\Omega$, it is deduced that:

$$\omega_+^0 P_+(\Omega)/P_\Omega(\Omega) \sim \hat{\omega}_+(\Omega) + O(\delta\Omega^2)$$

$$\omega_0^0 P_0(\Omega)/P_\Omega(\Omega) \sim \hat{\omega}_0(\Omega) + O(\delta\Omega^2)$$

$$\omega_-^0 P_-(\Omega)/P_\Omega(\Omega) \sim \hat{\omega}_-(\Omega) + O(\delta\Omega^2)$$

Thus, if the population is sufficiently large or dense, the direct mapping of fields is theoretically possible.

**SM4: Condition imposed by the conservation of genetic microstate frequencies when the minimalist model (Fisher's fields) is considered**

Let us write '$1/\Delta(\Omega)$'as '$\Delta^{-1}(\Omega)$' for concise notations and assume that the population is large and dense enough to assimilate '$\Delta^{-1}(\Omega)$' to the phenotype distribution density function (see SM1).

Recalling the fields as defined in SM2 (see Eq.SM2-1). One sets $\hat{x} \stackrel{\text{def}}{=} s(\Omega - \Omega_m)$, where '$\Omega_m$' is defined in SM2 and chosen as the new origin of the phenotypic values. Consequently the new integration interval is $[\Omega_{1/N} - \Omega_m, \Omega_1 - \Omega_m]$. Provided that the population is large and dense enough (see SM1), using the asymptotical convergence in probability of '$\Delta^{-1}(\Omega)$', the integration interval can be extended to $]-\infty, +\infty[$. Note that this extension is artificial, but mathematically useful.

As a result, the conservation of genetic microstate frequencies (Eq.8 main text) when the minimalist model is employed (see SM2 or Eqs.12 main text) can be rewritten as:

$$\int_{-\infty}^{+\infty} \frac{sh(\hat{x}_0-\hat{x})}{\alpha_0+ch(\hat{x}_0-\hat{x})} \Delta^{-1}\left(\frac{\hat{x}}{s}\right)\frac{d\hat{x}}{s} \sim \frac{sh(\hat{x}_0)}{\alpha_0+ch(\hat{x}_0)}$$

Recall that '$\alpha_0$' is constant. This integral can be determined in a complex map by using Cauchy's integral theorem. To use this theorem, one notes by $z = a + ib$ any complex number where 'a' and 'b' are real numbers and $i^2 = -1$; and rewrites the integral in the form:

$$\oint_C \frac{sh(\hat{x}_0-\hat{z})}{\alpha_0+ch(\hat{x}_0-\hat{z})} \Delta^{-1}\left(\frac{\hat{z}}{s}\right)\frac{d\hat{z}}{s} \sim \frac{sh(\hat{x}_0)}{\alpha_0+c\ \ (\hat{x}_0)}$$

where '$C$' is a closed contour to be determined.

Cauchy's integral theorem states that the latter integral (i.e. the left-end side) can be re-expressed as a discreet sum of residues, provided that the integrand is holomorphic within the area delineated by the closed contour '$C$', namely that the contour must be drawn in such a way that the poles are excluded.

The contour chosen is a semicircle extending along the positive values of the imaginary axis and whose diameter is formed by the real axis. Such a contour can be parametrized by the complex equation: $Re^{i\varphi}$ where ‘R’ is the radius of the semi-circle and $\varphi \in [0;\pi]$ the polar angle. Using Jordan’s lemma, the curved part of the contour can be demonstrated to be a ghost contour without difficulty, owing to the rapid asymptotic convergence in the probability of ‘$\Delta^{-1}$’.

The poles are defined when the denominator of the integrand is null and is therefore a solution of $\mathrm{ch}(\hat{x}_0 - \hat{z}) = -\alpha_0$. Using the exponential expression of the hyperbolic function, we find that the poles are therefore a solution of $e^{2(\hat{x}_0-\hat{z})} + 2\alpha_0 e^{(\hat{x}_0-\hat{z})} + 1 = 0$. This quadratic equation can be solved with the following solutions:

$$e^{(\hat{x}_0-(\hat{z}_-)_n)} = -\left(\alpha_0 + \sqrt{{\alpha_0}^2 - 1}\right) = e^{-(2n+1)i\pi} e^{\ln\left(\alpha_0+\sqrt{{\alpha_0}^2-1}\right)}$$

$$e^{(\hat{x}_0-(\hat{z}_+)_n)} = -\left(\alpha_0 - \sqrt{{\alpha_0}^2 - 1}\right) = e^{-(2n+1)i\pi} e^{\ln\left(\alpha_0-\sqrt{{\alpha_0}^2-1}\right)}$$

where ‘n’ is a relative number.

Therefore, there are two single roots when $\alpha_0 \neq 1$ and one double root when $\alpha_0 = 1$. One recalls that $\alpha_0 = 1$ corresponds to the Hardy-Weinberg law.

In this context, there is a need to discuss two cases: $\alpha_0 \neq 1$ and $\alpha_0 = 1$.

**Treatment for the case $\alpha_0 = 1$:**

This case is simple to handle, and provides the main tools, ideas, and simplifications used.

The double root, imposes a set of poles determined by:

$(\hat{z}_0)_n = \hat{x}_0 + (2n + 1)i\pi$, where ‘n’ is a relative number. In this case, the residues for the poles are calculated for any value of ‘n’ using the limit

$$\mathrm{Res}((\hat{z}_0)_n) = \lim_{\varepsilon \to 0} \oint_{C_\varepsilon} \frac{\mathrm{sh}\left(\hat{x}_0-(\hat{z}_0)_n-\varepsilon e^{i\beta}\right)}{1+\mathrm{ch}\left(\hat{x}_0-(\hat{z}_0)_n-\varepsilon e^{i\beta}\right)} \Delta^{-1}\left(\frac{(\hat{z}_0)_n+\varepsilon e^{i\beta}}{s}\right) \frac{\varepsilon e^{i\beta} i d\beta}{s}$$

Where ‘$C_\varepsilon$’ is an anticlock-wise contour surrounding any given pole.

One deduces then:

$$\mathrm{Res}((\hat{z}_0)_n) = \lim_{\varepsilon\to 0} \oint_{C_\varepsilon} \frac{\mathrm{sh}\left((2n+\ )i\pi - \varepsilon e^{i\beta}\right)}{1+\mathrm{ch}\left((2n+1)i\pi - \varepsilon e^{i\beta}\right)} \Delta^{-1}\left(\frac{\hat{x}_0+(2n+1)i\pi+\varepsilon e^{i\beta}}{s}\right)\frac{\varepsilon e^{i\beta} i d\beta}{s}$$

Because the hyperbolic functions are 2πi-periodic, the residues can be rewritten in the following form:

$$\mathrm{Res}((\hat{z}_0)_n) = \lim_{\varepsilon\to 0} \oint_{C_\varepsilon} \frac{\mathrm{sh}\left(i\pi - \varepsilon e^{i\beta}\right)}{1+\mathrm{ch}\left(i\pi - \varepsilon e^{i\beta}\right)} \Delta^{-1}\left(\frac{\hat{x}_0+(2n+1)i\pi+\varepsilon e^{i\beta}}{s}\right)\frac{\varepsilon e^{i\beta} i d}{s}$$

Considering $\varepsilon \to 0$ the chain rule method applied to the hyperbolic functions in the integrand yields:

$$\mathrm{sh}\left(i\pi - \varepsilon e^{i\beta}\right) \sim \mathrm{sh}(i\pi) - \mathrm{ch}(i\pi)\varepsilon e^{i\beta} + \frac{1}{2}\mathrm{sh}(i\pi)\left(\varepsilon e^{i\beta}\right)^2 + O(\varepsilon^3)$$

$$\mathrm{ch}\left(i\pi - \varepsilon e^{i\beta}\right) \sim \mathrm{ch}(i\pi) - \mathrm{sh}(i\pi)\varepsilon e^{i\beta} + \frac{1}{2}\mathrm{ch}(i\pi)\left(\varepsilon e^{i\beta}\right)^2 + O(\varepsilon^3)$$

As $\mathrm{sh}(i\pi) = 0$ and $\mathrm{ch}(i\pi) = -1$ one deduces:

$$\mathrm{sh}\left(i\pi - \varepsilon e^{i\beta}\right) \sim +\varepsilon e^{i\beta} + O(\varepsilon^3)$$

$$\mathrm{ch}\left(i\pi - \varepsilon e^{i\beta}\right) \sim -1 - \frac{1}{2}\left(\varepsilon e^{i\beta}\right)^2 + O(\varepsilon^3)$$

As a result:

$$\mathrm{Res}((\hat{z}_0)_n) = \lim_{\varepsilon\to 0} \oint_{C_\varepsilon} \frac{\mathrm{sh}\left(i\pi - \varepsilon e^{i\beta}\right)}{1+\mathrm{ch}\left(i\pi - \varepsilon e^{i\beta}\right)} \Delta^{-1}\left(\frac{\hat{x}_0+(2n+1)i\pi+\varepsilon e^{i\beta}}{s}\right)\frac{\varepsilon e^{i\beta} i d\beta}{s} \sim 4i\pi\Delta^{-1}\left(\frac{\hat{x}_0+(2n+1)i\pi}{s}\right)\frac{1}{s}$$

Finally, one finds that Eq.8 (main text), together with Fisher fields, imposes

$$\sum_{n=0}^{+\infty} 4i\pi\Delta^{-1}\left(\frac{\hat{x}_0+(2n+1)i\pi}{s}\right)\frac{1}{s} = \frac{\mathrm{sh}(\hat{x}_0)}{1+\mathrm{ch}(\hat{x}_0)}$$

In order to simplify the summation over ‘n’, let us recall that that ‘$\hat{x}_0$’ is a silent variable and transform: $\hat{x}_0 \to \hat{y}_0 \stackrel{\text{def}}{=} \hat{x}_0 + 2i\pi$; such as to rewrite the summation as:

$$\sum_{n=0}^{+\infty} 4i\pi\Delta^{-1}\left(\frac{\hat{y}_0+(2(n-1)+1)i\pi}{s}\right)\frac{1}{s} = \frac{\mathrm{sh}(\hat{y}_0-2i\pi)}{1+\mathrm{ch}(\hat{y}_0-2i\pi)} = \frac{\mathrm{sh}(\hat{y}_0)}{1+\mathrm{ch}(\hat{y}_0)}$$

By changing the starting value of ‘n’ upon which the summation starts, we deduce

$$\sum_{n=-1}^{+\infty} 4i\pi\Delta^{-1}\left(\frac{\hat{y}_0+(2n+1)i\pi}{s}\right)\frac{1}{s} = \frac{\mathrm{sh}(\hat{y}_0)}{1+\mathrm{ch}(\hat{y}_0)} = \sum_{n=0}^{+\infty} 4i\pi\Delta^{-1}\left(\frac{\hat{y}_0+(2n+1)i\pi}{s}\right)\frac{1}{s}$$

As a result, the conservation of genetic microstates when Fisher's fields are employed and when Hardy-Weinberg law is valid is $\Delta^{-1}\left(\frac{\hat{y}_0 - i\pi}{s}\right) = 0$, or equivalently and with our notations: $\Delta^{-1}\left(\frac{\hat{x}_0 + i\pi}{s}\right) = 0$

Thus, provided that the phenotype distribution density function is given, the latter relation provides the conditions linked to the validity of Eq.8 when Fisher's fields are employed in a context where Hardy-Weinberg law is verified.

**Treatment for the case $\alpha_0 \neq 1$:**

Let us assume $\alpha_0 > 1$ for the moment, as we shall see that specific symmetries will arise, which will allow us to deduce the case when $\alpha_0 < 1$.

Assuming $\alpha_0$ >1 or equivalently $1 + \Delta\omega_0{}^2 \geq 2\omega_0$, one deduces that the poles are given by

$$(\hat{z}_-)_n = \hat{x}_0 - \ln\left(\alpha_0 + \sqrt{\alpha_0{}^2 - 1}\right) + (2n+1)i\pi \qquad \text{(SM4-1a)}$$

$$(\hat{z}_+)_n = \hat{x}_0 - \ln\left(\alpha_0 - \sqrt{\alpha_0{}^2 - 1}\right) + (2n+1)i\pi \qquad \text{(SM4-1b)}$$

By posing $\theta_- = \ln\left(\alpha_0 - \sqrt{\alpha_0{}^2 - 1}\right)$ and $\theta_+ = \ln\left(\alpha_0 + \sqrt{\alpha_0{}^2 - 1}\right)$, one notes that $\theta_+ + \theta_- = \ln(1) = 0$ and $\theta_+ - \theta_- = \ln\left(\frac{\alpha_0 + \sqrt{\alpha_0{}^2-1}}{\alpha_0 - \sqrt{\alpha_0{}^2-1}}\right) = 2\mathrm{ath}\left(\sqrt{1 - \frac{1}{\alpha_0^2}}\right)$. As a result, $(\theta_+ + \theta_-) + (\theta_+ - \theta_-) = 2\theta_+ = 2\mathrm{ath}\left(\sqrt{1 - \frac{1}{\alpha_0^2}}\right)$ and $(\theta_+ + \theta_-) - (\theta_+ - \theta_-) = 2\theta_- = -2\mathrm{ath}\left(\sqrt{1 - \frac{1}{\alpha_0^2}}\right)$.

Then by defining $\theta_+ = -\theta_- \stackrel{\text{def}}{=} \theta_0$ it follows $\mathrm{th}(\theta_0) = \sqrt{1 - \frac{1}{\alpha_0^2}}$ or equivalently $\alpha_0^2 = 1/\left(1 - \mathrm{th}^2(\theta_0)\right)$. Because $\mathrm{ch}^2(\theta_0) = 1/\left(1 - \mathrm{th}^2(\theta_0)\right)$; we can also define $\alpha_0 = \mathrm{ch}(\theta_0)$.

One deduces then, that Eq.SM4-1a and Eq.SM4-1b can be simplified when $\alpha_0 \geq 1$ and rewritten as:

$$(\hat{z}_-)_n = \hat{x}_0 - \theta_0 + (2n+1)i\pi \qquad \text{(SM4-2a)}$$

$$(\hat{z}_+)_n = \hat{x}_0 + \theta_0 + (2n+1)i\pi \qquad \text{(SM4-2b)}$$

Consider now that, $\alpha_0 < 1$ or equivalently $1 + \Delta\omega_0{}^2 < 2\omega_0$. The poles can be rewritten as:

$$(\hat{z}_-)_n = \hat{x}_0 - \ln\left(\alpha_0 + i\sqrt{1-\alpha_0{}^2}\right) + (2n+1)i\pi \qquad \text{(SM4-3a)}$$

$$(\hat{z}_+)_n = \hat{x}_0 - \ln\left(\alpha_0 - i\sqrt{1-\alpha_0{}^2}\right) + (2n+1)i\pi \qquad \text{(SM4-3b)}$$

The argument in the natural log can be simplified using Euler notation by posing $\alpha_0 \pm i\sqrt{1-\alpha_0{}^2} = e^{\pm i\theta_0}$ with $\tan(\theta_0) = \sqrt{\frac{1}{\alpha_0^2} - 1}$ or equivalently, $\alpha_0^2 = 1/\left(1+\tan^2(\theta_0)\right)$.

Because $\cos^2(\theta_0) = 1/\left(1+\tan^2(\theta_0)\right)$, the latter relation allows the definition of $\alpha_0 = \cos(\theta_0) = \mathrm{ch}(i\theta_0)$.

One deduces then, that Eq.SM-3a and Eq.SM-3b can be simplified when $\alpha_0 < 1$ and rewritten as:

$$(\hat{z}_-)_n = \hat{x}_0 - i\theta_0 + (2n+1)i\pi \qquad \text{(SM4-4a)}$$

$$(\hat{z}_+)_n = \hat{x}_0 + i\theta_0 + (2n+1)i\pi \qquad \text{(SM4-4b)}$$

The only difference between $\alpha_0 > 1$ and $\alpha_0 < 1$ lies in the transformation $\theta_0 \to i\theta_0$. As a result, there is a fundamental difference in the solutions obtained between the two aforementioned cases linked to the occurrence of semi-periodic solutions connected to the microstate frequencies in the complex map.

To provide a more concise notation one shall replace '$\theta_0$' or '$i\theta_0$' by '$\epsilon\theta_0$' with the understanding that, $\epsilon = 1$ if $\alpha_0 > 1$ or, $\epsilon = i$ if $\alpha_0 < 1$.

The determination of residues is then performed by calculating:

$$\mathrm{Res}((\hat{z}_+)_n) = \lim_{\varepsilon\to 0} \oint_{C_\varepsilon} \frac{\mathrm{sh}\left(-\epsilon\theta_0 - (2n+1)i\pi - \varepsilon e^{i\beta}\right)}{\mathrm{ch}(\epsilon\theta_0) + \mathrm{ch}\left(-\epsilon\theta_0 - (2n+1)i\pi - \varepsilon e^{i\beta}\right)} \Delta^{-1}\left(\frac{\hat{x}_0 + \epsilon\theta_0 + (2n+1)i\pi + \varepsilon e^{i\beta}}{s}\right) \frac{\varepsilon e^{i\beta} i d\beta}{s}$$

$$\mathrm{Res}((\hat{z}_-)_n) = \lim_{\varepsilon\to 0} \oint_{C_\varepsilon} \frac{\mathrm{sh}\left(+\epsilon\theta_0 - (2n+1)i\pi - \varepsilon e^{i\beta}\right)}{\mathrm{ch}(\epsilon\theta_0) + \mathrm{ch}\left(+\epsilon\theta_0 - (2n+1)i\pi - \varepsilon e^{i\beta}\right)} \Delta^{-1}\left(\frac{\hat{x}_0 - \epsilon\theta_0 + (2n+1)i\pi + \varepsilon e^{i\beta}}{s}\right) \frac{\varepsilon e^{i\beta} i d\beta}{s}$$

Where '$C_\varepsilon$' is an anticlock-wise contour surrounding any given pole.

As hyperbolic functions are 2iπ-periodic the integrands can be simplified as:

$$\mathrm{Res}((\hat{z}_{+})_{n}) = \lim_{\varepsilon\to 0}\oint_{C_\varepsilon} \frac{-\mathrm{s}\ \ (\epsilon\theta_0+i\pi+\varepsilon e^{i\beta})}{\mathrm{ch}(\epsilon\theta_0)+\mathrm{ch}(\epsilon\theta_0+i\pi+\varepsilon e^{i\beta})}\Delta^{-1}\left(\frac{\hat{x}_0+\epsilon\theta_0+(2n+1)i\pi+\varepsilon e^{i\beta}}{s}\right)\frac{\varepsilon e^{i\beta} i d\beta}{s}$$

$$\mathrm{Res}((\hat{z}_{-})_{n}) = \lim_{\varepsilon\to 0}\oint_{C_\varepsilon} \frac{\mathrm{sh}(+\epsilon\theta_0-i\pi-\varepsilon e^{i\beta})}{\mathrm{ch}(\epsilon\theta_0)+\mathrm{c}\ \ (+\epsilon\theta_0-i\pi-\varepsilon e^{i\beta})}\Delta^{-1}\left(\frac{\hat{x}_0-\epsilon\theta_0+(2n+1)i\pi+\varepsilon e^{i\beta}}{s}\right)\frac{\varepsilon e^{i\beta} i d\beta}{s}$$

The hyperbolic function in the integrands can be further developed and approximated by considering the limit $\varepsilon \to 0$:

$$\mathrm{sh}(\epsilon\theta_0 + i\pi + \varepsilon e^{i\beta}) \sim -\mathrm{sh}(\epsilon\theta_0) - \mathrm{ch}(\epsilon\theta_0)\varepsilon e^{i\beta} + O(\varepsilon^2)$$

$$\mathrm{ch}(\epsilon\theta_0 + i\pi + \varepsilon e^{i\beta}) \sim -\mathrm{ch}(\epsilon\theta_0) - \mathrm{sh}(\epsilon\theta_0)\varepsilon e^{i\beta} + O(\varepsilon^2)$$

$$\mathrm{sh}(\epsilon\theta_0 - i\pi - \varepsilon e^{i\beta}) \sim -\mathrm{sh}(\epsilon\theta_0) + \mathrm{ch}(\epsilon\theta_0)\varepsilon e^{i\beta} + O(\varepsilon^2)$$

$$\mathrm{ch}(\epsilon\theta_0 - i\pi - \varepsilon e^{i\beta}) \sim -\mathrm{ch}(\epsilon\theta_0) + \mathrm{sh}(\epsilon\theta_0)\varepsilon e^{i\beta} + O(\varepsilon^2)$$

One deduces then:

$$\mathrm{Res}((\hat{z}_{+})_{n}) = \frac{2i\pi}{s}\Delta^{-1}\left(\frac{\hat{x}_0+\epsilon\theta_0+(2n+1)i\pi}{s}\right)$$

$$\mathrm{Res}((\hat{z}_{-})_{n}) = \frac{2i\pi}{s}\Delta^{-1}\left(\frac{\hat{x}_0-\epsilon\theta_0+(2n+1)i\pi}{s}\right)$$

As a result, the conservation of genetic microstate frequencies (Eq.8 main text) when Fisher's fields are employed can be rewritten, as:

$$\sum_{n=0}^{+\infty}\Delta^{-1}\left(\frac{\hat{x}_0}{s}+\frac{\epsilon\theta_0+(2n+1)i\pi}{s}\right)+\Delta^{-1}\left(\frac{\hat{x}_0}{s}-\frac{\epsilon\theta_0-(2n+1)i\pi}{s}\right)\sim\frac{s}{2\pi}\frac{i\cdot \mathrm{sh}(\hat{x}_0)}{\mathrm{ch}(\epsilon\theta_0)+\mathrm{ch}(\hat{x}_0)} \qquad \text{(SM4-5)}$$

From Eq.SM4-5, symmetries are clearly visible.

Recalling that with the notations used: $\alpha_0 = \mathrm{ch}(\epsilon\theta_0) = (1-\omega_0)\mathrm{ch}(\hat{x}_0)/\omega_0$, and that the hyperbolic sine and cosine functions are 2iπ-periodic, one rewrite '$\alpha_0$' under the form: $\alpha_0 = \mathrm{ch}(\epsilon\theta_0) = (1-\omega_0)\mathrm{ch}(\hat{x}_0 + 2i\pi)/\omega_0$.

The latter result shows that '$\epsilon\theta_0$' is invariant upon the transformation: $\hat{x}_0 \to \hat{y}_0 \stackrel{\text{def}}{=} \hat{x}_0 + 2i\pi$.

Let us then rewrite Eq.SM4-5 as:

$$\sum_{n=0}^{+\infty}\Delta^{-1}\left(\frac{\hat{y}_0}{s}+\frac{\epsilon\theta_0+(2(n-1)+1)i\pi}{s}\right)+\Delta^{-1}\left(\frac{\hat{y}_0}{s}-\frac{\epsilon\theta_0-(2(n-1)+1)i\pi}{s}\right)\sim\frac{s}{2\pi}\frac{i\cdot \mathrm{sh}(\hat{y}_0-2i\pi)}{\mathrm{ch}(\epsilon\theta_0)+\mathrm{ch}(\hat{y}_0-2i\pi)}$$

The starting value of 'n' can then be changed transforming the sum as:

$$\sum_{n=-1}^{+\infty} \Delta^{-1}\left(\frac{\hat{y}_0}{s}+\frac{\epsilon\theta_0+(2n+1)i\pi}{s}\right)+\Delta^{-1}\left(\frac{\hat{y}_0}{s}-\frac{\epsilon\theta_0-(2n+1)i\pi}{s}\right)\sim\frac{s}{2\pi}\frac{i\cdot sh(\hat{y}_0)}{ch(\epsilon\theta_0)+c\ \ (\hat{y}_0)}$$

Making use of Eq.SM4-5 one deduces then:

$$\Delta^{-1}\left(\frac{\hat{y}_0}{s}+\frac{\epsilon\theta_0-i\pi}{s}\right)+\Delta^{-1}\left(\frac{\hat{y}_0}{s}-\frac{\epsilon\theta_0+i\pi}{s}\right)\sim 0$$

Finally, by reintroducing the variable '$\hat{x}_0$' one finds finally:

$$\Delta^{-1}\left(\frac{\hat{x}_0+i\pi}{s}+\frac{\epsilon\theta_0}{s}\right)+\Delta^{-1}\left(\frac{\hat{x}_0+i\pi}{s}-\frac{\epsilon\theta_0}{s}\right)\sim 0 \qquad \text{(SM4-6)}$$

The latter relation provides the constraint linked to Eq.8 (main text) that arises between the genetic microstate frequencies characterising a genome position and the phenotype distribution density function when Fisher's fields are employed. If the phenotype distribution density is known, then its mathematical expression can be included in Eq.SM4-6 to deduce how the variables linked to microstate frequencies must be related.

### SM5: Code for the simulations

### SM5-a: Study Power

```
%GWASpower.m
%Circ 9 March 2022
%script to calculate power of a standard GWAS using formulae from
Delongchamp R, Faramawi MF, Feingold E, Chung D, Abouelenein S.
%The Association between SNPs and a Quantitative Trait: Power
Calculation. Eur J Environ Public Health. 2018;2(2):10. doi:
10.20897/ejeph/3925


%non-centrality parameter (NCP) in an additive genetic model

%set the allele effect and frequency and the sample size (number of
individuals)
%==========================
alleff=[0.00001 0.0001 0.0005 0.001 0.005 0.01 0.02 0.05 0.06 0.08 0.1
0.15 0.2 0.25 0.3 0.5 0.8 1];    %allele effect

freq_A1=0.8;                       %allele frequency
n=[1000 10000 100000 1000000];     %sample size - number of individuals
%==========================
%calculate the genotype frequencies
freq_A2=1-freq_A1;
freq_A1A1=(freq_A1)^2;
freq_A1A2=2*freq_A1*freq_A2;
freq_A2A2=(freq_A2)^2;

neff=length(alleff);
nN=length(n);
output=zeros(neff,nN);

hold
for i=1:nN;
    for j=1:neff;
    %calculate NCP
    freq_factor=1*freq_A1A1+0*freq_A1A2+1*freq_A2A2;
    ncp=n(i)*(alleff(j))^2*freq_factor;  %non-centrality parameter
    %set type I error (significance threshold)
    a=0.00000005  ;     %5 x 10-8 is standard threshold suggested by
Risch and Merikangas (1996), Panagiotou OA , Ioannidis JPA, (2012)
    %for the Genome-Wide Significance Project 2012. What should the
genome-wide significance threshold be? Empirical replication of
borderline genetic associations.     %Int J Epidemiol. 41:273–286.
doi:10.1093/ije/dyr178
    p=0;                %other covariates
    d=2;                %degrees of freedom - genotypes
    v=n(i)-p-d;            %degress of freedom - error
    c=finv(1-a,d,v);
    %calculate power
    power=1-ncfcdf(c,d,v,ncp);
    output(j,i)=power;
    end
    plot(alleff,output(:,i),'o');
end
```

**SM5-b: Genotype-phenotype-position**

```
%gensimHeight.m
%Circ 9 March 2022
%MATLAB script generating simulated data for Rauch et al.

&======README comments ==========

%This script simulates genotypes with human height as the phenotype.
Mean and variance of human height (mixed males and females) are derived
from GTEx project data (www.gtexportal.org).  Genotypes are simulated
according to exact Hardy-Weinberg Equilibrium proportions expected e.g.
if p=0.5 proportions of -1,0,+1 genotypes, %where -1 individuals are
the tallest and +1 individuals the smallest, are p2, 2pq, q2
%The script can be run over multiple values of gene effect (a)
automatically.  Different values of allele frequency (p) need to be
inserted manually.  Saved output is theta diff stats - max value, posn,
phenotype and standard deviations.  Also vector
%of theta diff and phenotype and natural log of theta diff for each set
of parameter values (a and p)

%=======beginning of code =======

%maxTheta_stats=[];

for k=1:11  %LOOP OVER NUMBER OF ALLELE EFFECTS TO BE SIMULATED.  k=1
to number of elements in vector alleff (see below).  The allele
frequency freq_A1 or p            %needs to be manually inserted below
for each allele frequency required.  Each set of allele effects is run
for a single allele frequency.

%=====SET PARAMETERS: allele frequencies,variance of the QTL,number of
individuals, number of replicates to run simulation for
nind=100;             %number of individuals to simulate
nrep=1000;           %number of simulated replicates
freq_A1=0.9;         %allele frequency
freq_A2=1-freq_A1;   %frequency of allele 2 is 1-frequency of allele 1
alleff=[1  0.8  0.5  0.3  0.15  0.1 0.05 0.01  0.005 0.001  0.0005];
%allele effect

%=======define gene effect, dominance, genetic variance and genotypic
values ===
a=alleff(k);                   %setting the value of a in phenotypic sd
d=0;     %defining the inter-allelic interaction or dominance
var_QTL=(2*freq_A1*freq_A2)*(a+d*(freq_A2-freq_A1))^2;
%QTL allele substitution effects and genetic variance
%See Falconer pg 117 and pg 129
%using notation from Falconer pg 112
   %A1A1 has genotypic value +a, A2A2 has genotypic value -a
   %and A1A2 has genotypic value d
   gval_A1A1=a;
   gval_A2A2=-a;
   gval_A1A2=d;

%==========calculate genotype frequencies
freq_A1A1=(freq_A1)^2;
freq_A1A2=2*freq_A1*freq_A2;
freq_A2A2=(freq_A2)^2;
```

```
%creating arrays to save data over each replicate
data_geno=[];
data_pheno=[];
ordered_geno=[];
ordered_pheno=[];
theta_diff=[];
theta_ordered=[];
theta_random=[];
maxTheta=[];
maxTheta_posn=[];
maxTheta_pheno=[];
sum_p2q2=[];
diff_p2q2=[];
het_freq=[];    %this is 2pq
estimated_a=[];
phenotype_stats=[];


for j=1:nrep            %LOOP FOR SIMULATION REPLICATES
%==========set up array of individual genotypes and phenotypes
genotype=zeros(nind,1);
phenotype=zeros(nind,1);
posn=[1:nind]';

%==genotypes now generated for exact genotype frequencies
nA1A1=int32(freq_A1A1*nind);        %stored as an integer value
nA1A2=int32(freq_A1A2*nind);
nA2A2=int32(freq_A2A2*nind);
total_ind=nA1A1+nA1A2+nA2A2;
total_gen=[nA1A1; nA1A2; nA2A2];
if total_ind>nind
    subtract=total_ind-nind;
    [mostfreq,idx]=max(total_gen);
    if idx==1
        nA1A1=nA1A1-subtract;
    elseif idx==2
        nA1A2=nA1A2-subtract;
    elseif idx==3
        nA2A2=nA2A2-subtract;
    end
end

%A1A1 individuals
for i=1:nA1A1
    genostate=-1;     %genotype A1A1
    genotype(i)=genostate;
    %generating environmental effect
    environ=normrnd(0,sqrt(1-var_QTL));
    phenotype(i)=gval_A1A1+environ;
end
for i=nA1A1+1:nA1A1+nA1A2
    genostate=0;      %genotype A1A2
    genotype(i)=genostate;
    %generating environmental effect
    environ=normrnd(0,sqrt(1-var_QTL));
    phenotype(i)=gval_A1A2+environ;
end
```

```
for i=nA1A1+nA1A2+1:nA1A1+nA1A2+nA2A2
    genostate=1;     %genotype A2A2
    genotype(i)=genostate;
    %generating environmental effect
    environ=normrnd(0,sqrt(1-var_QTL));
    phenotype(i)=gval_A2A2+environ;
end

%translating phenotype to realistic values
height_mean=68.17206;      %overall mean height of 832 GTEx donors in
inches (males and females)
height_sd=4.030067;         %sd of height (males and females)
phenotype=(phenotype*height_sd)+height_mean;

%randomise genotype order - as they were generated by groups of
genotype
random_idx=randperm(nA1A1+nA1A2+nA2A2)';
genotype=genotype(random_idx);
phenotype=phenotype(random_idx);   %phenotype still needs to be linked
to correct genotype

%saving genotype and phenotype to arrays for further analysis
data_geno=[data_geno genotype];
data_pheno=[data_pheno phenotype];

%theta diff
S=cumsum(genotype);   %theta random
[b,pheno_idx]=sort(phenotype);   %ordering by phenotype
S_pheno=cumsum(genotype(pheno_idx));  %theta ordered
S_diff=S_pheno-S;    %theta diff

%find max theta, its posn and phenotype value - the max may cover
%several positions so am taking the mean over those positions
max_theta_idx=find(S_diff==max(S_diff));  %finding the maximum value(s)
of theta diff
max_theta=mean(S_diff(max_theta_idx));        %value of max theta diff
max_theta_posn=mean(posn(max_theta_idx));     %calculating mean
position for max theta diff
max_theta_pheno=mean(b(max_theta_idx));       %calculating mean
phenotype for max theta diff

%save data over each replicate
ordered_geno=[ordered_geno genotype(pheno_idx)];
ordered_pheno=[ordered_pheno phenotype(pheno_idx)];
theta_ordered=[theta_ordered S_pheno];
theta_random=[theta_random S];
theta_diff=[theta_diff S_diff];
maxTheta=[maxTheta max_theta];
maxTheta_posn=[maxTheta_posn max_theta_posn];
maxTheta_pheno=[maxTheta_pheno max_theta_pheno];

%find the three genotypes/microstates - to use for estimating gene
effect and colour coding graphs
idx_plus_one=find(genotype==1);
idx_minus_one=find(genotype==-1);
idx_zero=find(genotype==0);
```

```
%estimating the gene effect a from the difference between homozygote
means
mean_p2=mean(phenotype(idx_minus_one));
std_p2=std(phenotype(idx_minus_one));
mean_q2=mean(phenotype(idx_plus_one));
std_q2=std(phenotype(idx_plus_one));
estimated_a=[estimated_a; (mean_q2-mean_p2)./2];   %calculating and
saving over replicates - this will be sd in units of measurement

end    %END OF LOOP OVER SIMULATION REPLICATES

%taking mean phenotype across replicates and normalising
mean_ordered_pheno=mean(ordered_pheno,2);
normal_ordered_pheno=mean_ordered_pheno./max(mean_ordered_pheno);
normal_posn=((1:nind)./nind)';
%calculating mean and sd of gene effect across replicates
mean_gene_effect=mean(estimated_a);
std_gene_effect=std(estimated_a);

%normalising mean and std of theta ordered
normal_tord_mean=(mean(theta_ordered,2))./nind;
normal_tord_std=(std(theta_ordered,0,2))./nind;
%normalising mean and std of theta random
normal_trand_mean=(mean(theta_random,2))./nind;
normal_trand_std=(std(theta_random,0,2))./nind;
%normalising mean and std of theta diff
normal_tdiff_mean=(mean(theta_diff,2))./nind;
normal_tdiff_std=(std(theta_diff,0,2))./nind;
%saving relevant data for each simulated scenario
normal_tdiff_data=[normal_posn mean_ordered_pheno normal_tdiff_mean
normal_tdiff_std];

%doing the same for the statistics on max theta
%for each simulated scenario there will be a single value of each
statistic
normal_maxTheta_mean=(mean(maxTheta,2))./nind;
normal_maxTheta_posn_mean=(mean(maxTheta_posn,2))./nind;
maxTheta_pheno_mean=(mean(maxTheta_pheno,2));
%standard deviations
normal_maxTheta_std=(std(maxTheta,0,2))./nind;
normal_maxTheta_posn_std=(std(maxTheta_posn,0,2))./nind;
maxTheta_pheno_std=(std(maxTheta_pheno,0,2));

%saving maxTheta statistics in a separate array
maxTheta_data=[a var_QTL normal_maxTheta_mean normal_maxTheta_std
normal_maxTheta_posn_mean normal_maxTheta_posn_std maxTheta_pheno_mean
maxTheta_pheno_std];

%phenotype and genotype(microstate) for last replicate of data
pheno_data=[genotype phenotype];

%====SAVING DATA========%
%writing data to Excel spreadsheet
sheet1=1;     %normalised mean and std theta diff - with normalised
position and phenotype
sheet2=2;     %max Theta stats
%sheet3=3;     %phenotype and genotype (microstate) data for last
replicate
```

```
if k<=7
shpos=['A' 'E' 'I' 'M' 'Q'  'U'  'Y'];   %Number of start points needs
to equal maximum value that k will attain
start=shpos(k);  shnum=string(1); sheet_posn=strcat(start,shnum);
elseif k>7
    shpos=['C'  'G'   'K'  'O'  'S'  'W'];
    new_shpos=strcat('A',shpos(k-7));
    start=new_shpos;  shnum=string(1); sheet_posn=strcat(start,shnum);
end
%defining range for maxTheta stats on sheet 2
rownum=k;    rgstart=string(rownum); range_start=strcat('A',rgstart);
range_end=strcat('H',rgstart);
range=strcat(range_start,':',range_end);
%defining sheet position for sheet 3
%shpos_sheet3=['A'  'C'  'E'  'G' 'I' 'K' 'M' 'O'  'Q'];
%sheet3_start=shpos_sheet3(k); sheet3_shnum=string(1);
sheet3_posn=strcat(sheet3_start,sheet3_shnum);

filename ='thetaOrdered_data.xlsx';    %normalised data written out to
the thetaOrdered_data.xlsx file
xlswrite(filename,normal_tdiff_data,sheet1,sheet_posn);
xlswrite(filename,maxTheta_data,sheet2,range);
%xlswrite(filename,pheno_data,sheet3,sheet3_posn);


end   %END OF LOOP OVER DIFFERENT ALLELE EFFECTS (k)

%=====PLOTTING DATA =======
%theta diff by phenotype, mean over all replicates
figure(1)
plot(mean_ordered_pheno,normal_tdiff_mean,'ro');
```

**SM6: Convergence and Self-consistency of GIFT**

**SM6.a: GIFT's convergence: Master equation for the genetic paths difference**

Here, we assume that the genetic paths are normalised by population size. To determine the convergence of GIFT, we return to the genetic path defined in the space of positions, and consider a string of genetic microstates for one particular genome position once the individuals have been ranked as a function of their phenotype value. As defined by Eq.1, the cumulative sum of the genetic microstates in the space of positions is '$\theta_{\hat{\imath}}$'.

Then we define the probability '$P(\hat{\imath},\theta)$' that the genetic path is of length '$\theta$' at the position '$\hat{\imath}$'. Treating '$\theta$' as a random variable for which the set of values it can take is constrained by the presence probability of genetic microstates, will allow us to determine the variance of the genetic paths difference.

We assume that the fields are defined such that $\omega_+(\hat{\imath})$, $\omega_0(\hat{\imath})$ and $\omega_-(\hat{\imath})$ are known for every position. Recalling that between two consecutive positions, say '$\hat{\imath}$' and '$\hat{\imath}+1/N$', the increment that '$\theta$' can take is either '+1/N', '0/N' or '-1/N'. Whether '$\theta$' is incremented by '+1/N', '0/N' or '-1/N' is determined by the presence probability of genetic microstates '+1/N', '0/N' or '-1/N' at the position '$\hat{\imath}+1/N$'. In this context, the probability '$P(\hat{\imath},\theta)$' can be expressed in the form of a recurring relation between two consecutive positions:

$$P(\hat{\imath}+1/N,\theta) = P(\hat{\imath},\theta-1/N)\omega_+(\hat{\imath}+1/N) + P(\hat{\imath},\theta)\omega_0(\hat{\imath}+1/N) + P(\hat{\imath},\theta+1/N)\omega_-(\hat{\imath}+1/N)$$

Where '$\omega_+$', '$\omega_0$' and '$\omega_-$' are the presence probabilities of genetic microstates '+1', '0' and '-1' at the position '$\hat{\imath}+1/N$'. Because $\omega_+(\hat{\imath})+\omega_0(\hat{\imath})+\omega_-(\hat{\imath}) = 1$ for any value of '$\hat{\imath}$', we deduce the following by using the continuum limit:

$$\partial_{\hat{\imath}}P(\hat{\imath},\theta) \sim -\Delta\omega\left(\hat{\imath}+\frac{1}{N}\right)\partial_\theta P(\hat{\imath},\theta) + \frac{1-\omega_0\left(\hat{\imath}+\frac{1}{N}\right)}{2N}(\partial_\theta)^2 P(\hat{\imath},\theta) + O(N^{-2})$$

Where '$\partial$' designates partial derivatives and, $\Delta\omega(\hat{\imath}+1/N) \overset{\text{def}}{=} \omega_+(\hat{\imath}+1/N) - \omega_-(\hat{\imath}+1/N)$.

However, as mentioned above, the key information resides in the difference between phenotype-associated and default genetic paths. The default genetic path noted, $\theta_0$, is given by $\theta_0 \sim \Delta\omega_0 \hat{\imath}$ with $\Delta\omega_0 = \omega_+^0 - \omega_-^0$. One can then determine the probability, $P(\hat{\imath}, \Delta\theta)$, that the genetic paths difference, $\Delta\theta \stackrel{\text{def}}{=} \theta - \theta_0$, attains a certain value at the position '$\hat{\imath}$' as:

$$\partial_{\hat{\imath}} P(\hat{\imath}, \Delta\theta) \sim -\left[\Delta\omega\left(\hat{\imath} + \frac{1}{N}\right) - \Delta\omega_0\right] \partial_{\Delta\theta} P(\hat{\imath}, \Delta\theta) + \frac{1 - \omega_0\left(\hat{\imath} + \frac{1}{N}\right)}{2N} (\partial_{\Delta\theta})^2 P(\hat{\imath}, \Delta\theta)$$

In this context, assuming that the first individual located at position $1/N$ has a known genetic microstate given by $\theta_{\hat{\imath}=1/N}$, by defining the first value of the genetic path difference as $\Delta\theta_0 = \frac{\theta_{\hat{\imath}=1/N} - (\omega_+^0 - \omega_-^0)}{N}$, the Fourier transform can be used to resolve the master equation.

Let us note '$\hat{k}$' the conjugate variable of '$\Delta\theta$' in the reciprocal Fourier space. Recall that in this context any function or probability '$P(\Delta\theta, \hat{\imath})$' can be decomposed in the Fourier space as: $P(\Delta\theta, \hat{\imath}) = \int_{-\infty}^{+\infty} \frac{d\hat{k}}{2\pi} \hat{P}(\hat{k}, \hat{\imath}) \exp(z\hat{k}\Delta\theta)$; where '$\hat{P}(\hat{k}, \hat{\imath})$' is the Fourier transform of '$P(\Delta\theta, \hat{\imath})$' and '$z$' is the imaginary number i.e. $z^2 = -1$. The inverse Fourier transform is then: $\hat{P}(\hat{k}, \hat{\imath}) = \int_{-\infty}^{+\infty} d\Delta\theta\, P(\Delta\theta, \hat{\imath}) \exp(-z\hat{k}\Delta\theta)$. As a result, the master equation can be rewritten in the Fourier reciprocal space as

$$\frac{\partial \hat{P}(\hat{k}, \hat{\imath})}{\partial \hat{\imath}} \sim \left[-z\hat{k}\left[\Delta\omega\left(\hat{\imath} + \frac{1}{N}\right) - \Delta\omega_0\right] - \hat{k}^2 \frac{1 - \omega_0\left(\hat{\imath} + \frac{1}{N}\right)}{2N}\right] \hat{P}(\hat{k}, \hat{\imath})$$

Once integrated over the position the solution is then:

$$\hat{P}(\hat{k}, \hat{\imath}) \sim \hat{A}(\hat{k}) \exp\left[-z\hat{k}\Delta\Sigma_1(\hat{\imath}) - \hat{k}^2 \frac{\Sigma_2(\hat{\imath})}{2N}\right]$$

Where '$\hat{A}(\hat{k})$' is a constant that needs to be determined by an 'initial condition'. In our case the initial condition is $P(\Delta\theta, \hat{\imath} = 1/N) = \delta(\Delta\theta - \Delta\theta_0)$ where '$\delta$' is the Dirac distribution, this condition can be rewritten as:

$$\hat{P}\left(\hat{k}, \hat{\imath} = \frac{1}{N}\right) \sim \hat{A}(\hat{k}) \exp\left[-z\hat{k}\Delta\Sigma_1\left(\hat{\imath} = \frac{1}{N}\right) - \hat{k}^2 \frac{\Sigma_2\left(\hat{\imath} = \frac{1}{N}\right)}{2N}\right] = \int_{-\infty}^{+\infty} d\Delta\theta\, \delta(\Delta\theta -$$

$$\Delta\theta_0) \exp(-z\hat{k}\Delta\theta) = \exp(-z\hat{k}\Delta\theta_0).$$

As $\Delta\Sigma_1\left(\hat{\imath}=\frac{1}{N}\right)\sim 0$ and $\Sigma_2\left(\hat{\imath}=\frac{1}{N}\right)/2N\sim 0$, one finds: $\widehat{A}(\hat{k})\sim\exp(-z\hat{k}\Delta\theta_0)$. As a result:

$\widehat{P}(\hat{k},\hat{\imath})\sim\exp\left[-z\hat{k}(\Delta\theta_0-\Delta\Sigma_1(\hat{\imath}))-\hat{k}^2\frac{\Delta\Sigma_2(\hat{\imath})}{2N}\right]$

This relation can then be inserted into the Fourier transform of '$P(\Delta\theta,\hat{\imath})$' to provide

$P(\Delta\theta,\hat{\imath})\sim\frac{1}{\sqrt{2\pi\Sigma_2(\hat{\imath})/N}}\exp\left[-\frac{(\Delta\theta-\Delta\theta_0-\Delta\Sigma_1(\hat{\imath}))^2}{2\Sigma_2(\hat{\imath})/N}\right]$

Where, $\Delta\Sigma_1(\hat{\imath})\stackrel{\text{def}}{=}\int_{1/N}^{\hat{\imath}}\left[\Delta\omega\left(x+\frac{1}{N}\right)-\Delta\omega_0\right]dx$ and $\Sigma_2(\hat{\imath})\stackrel{\text{def}}{=}\int_{1/N}^{\hat{\imath}}\left[1-\omega_0\left(x+\frac{1}{N}\right)\right]dx$.

Using the symbol '$\langle\cdot\rangle$' to denote average values, it follows that the expected path is: $\langle\Delta\theta\rangle\sim\Delta\theta_0+\Delta\Sigma_1(\hat{\imath})$ with a variance given by $\langle(\Delta\theta-\Delta\theta_0-\Delta\Sigma_1(\hat{\imath}))^2\rangle\sim 2\Sigma_2(\hat{\imath})/N$; and both results can be rewritten informally as: $\Delta\theta\sim\Delta\theta_0+\Delta\Sigma_1(\hat{\imath})\pm\sqrt{2\Sigma_2(\hat{\imath})/N}$.

The key information from this development is that the genetic path is expected to converge rapidly as $\sim 1/\sqrt{N}$. One may also re-develop the relation above in the space of phenotypic values in place of positions. In this case, by assuming that 'N' is large enough, $\Delta\widehat{\Sigma}_1(\Omega)\sim\int_{\Omega_{1/N}}^{\Omega}[\Delta\widehat{\omega}(\Omega)-\Delta\omega_0]\frac{1}{\Delta(\Omega)}d\Omega+O(N^{-1})$ and $\widehat{\Sigma}_2(\Omega)\sim\int_{\Omega_{1/N}}^{\Omega}(1-\widehat{\omega}_0(\Omega))\frac{1}{\Delta(\Omega)}d\Omega+O(N^{-1})$. As $\Delta\theta_0\to 0$ with 'N' large, we deduce that:

$$P(\Delta\hat{\theta},\Omega)\sim\frac{1}{\sqrt{2\pi\widehat{\Sigma}_2(\Omega)/N}}\exp\left[-\frac{(\Delta\hat{\theta}-\Delta\widehat{\Sigma}_1(\Omega))^2}{2\widehat{\Sigma}_2(\Omega)/N}\right] \qquad \text{(SM6.a-1)}$$

With Eq.SM6.a-1 one can also consider the involvement of several paths. This exercise provides the overall self-consistency of the model (see SM6.b below).

**SM6.b: self-consistency of GIFT**

The genetic path difference is a function of phenotypic fields, which are in turn a function of phenotype values. Given that genome positions also participate in the creation of a set of phenotype values, the model must be self-consistent. Namely, a 'loop' regarding the set of phenotype values can be defined. As the phenotypic fields experienced by a genome position result from the effect of all genome positions involved in the phenotype considered, the fields also prescribe the impact of potential interactions between genes.

Therefore, concentrating on a set of genome positions that we shall note as '$\{\Delta\hat{\theta}_j\}$' with: $j = 1, \cdots, n$, where 'n' is the number of genome positions involved in the phenotype considered; the probability that the genetic paths difference of the $j^{th}$ genome position is '$\Delta\hat{\theta}_j$' for the phenotype value '$\Omega$' is given by Eq.SM6.a-1 under the form:

$$P_j(\Delta\hat{\theta}_j, \Omega) \sim \frac{1}{\sqrt{2\pi(\hat{\Sigma}_2)_j(\Omega)/N}} \exp\left[-\frac{\left(\Delta\hat{\theta}_j - (\Delta\hat{\Sigma}_1)_j(\Omega)\right)^2}{2(\hat{\Sigma}_2)_j(\Omega)/N}\right] \qquad \text{(SM6.b-1)}$$

Where, $(\Delta\hat{\Sigma}_1)_j(\Omega) \sim \int_{\Omega_{1/N}}^{\Omega} [(\Delta\hat{\omega})_j(\Omega) - (\Delta\omega_0)_j] \frac{1}{\Delta(\Omega)} d\Omega$, and $(\hat{\Sigma}_2)_j(\Omega) \sim \int_{\Omega_{1/N}}^{\Omega} \left(1 - (\hat{\omega}_0)_j(\Omega)\right) \frac{1}{\Delta(\Omega)} d\Omega$.

Let us consider a phenotype, as the genome positions specifically prescribe this phenotype, one can define by '$\hat{\Gamma}_\Omega$' the phenotypic path resulting from genome positions. Accordingly, we can then define a relation between '$\hat{\Gamma}_\Omega$' and the set $\{\Delta\hat{\theta}_j\}$ under the form: $\hat{\Gamma}_\Omega \stackrel{\text{def}}{=} F_\Omega(\{\Delta\hat{\theta}_j\})$, where '$F_\Omega$' is an unknown function prescribing the involvement of genome positions '$\{\Delta\hat{\theta}_j\}$' leading to the phenotypic path '$\hat{\Gamma}_\Omega$'. Thus, '$\hat{\Gamma}_\Omega$' is conditioned by $\{\Delta\hat{\theta}_j\}$ through '$F_\Omega$'. In this context, we can write the conditional probability for the occurrence of '$\hat{\Gamma}_\Omega$' under the form: $P(\hat{\Gamma}_\Omega/\{\Delta\hat{\theta}_j\}) = \delta\left(\hat{\Gamma}_\Omega - F_\Omega(\{\Delta\hat{\theta}_j\})\right)$, where '$\delta$' is the Dirac distribution.

Defining '$P(\hat{\Gamma}_\Omega)$' as the probability that '$\hat{\Gamma}_\Omega$' occurs, this probability is then written as

$$P(\hat{\Gamma}_\Omega) = \int \delta\left(\hat{\Gamma}_\Omega - F_\Omega(\{\Delta\hat{\theta}_j\})\right) \prod_{j=1}^{n} P_j(\Delta\hat{\theta}_j, \Omega) d\Delta\hat{\theta}_j \qquad \text{(SM6.b-2)}$$

Where the integration is performed over all the '$d\Delta\hat{\theta}_j$', with an integration interval $]-\infty, +\infty[$ and; '$\prod_{j=1}^{n}$' denotes the product of the different '$P_j(\Delta\hat{\theta}_j, \Omega) d\Delta\hat{\theta}_j$' for $j = 1, \cdots, n$.

Noting $\widetilde{P}(k) \stackrel{\text{def}}{=} \int e^{ik\hat{\Gamma}_\Omega} P(\hat{\Gamma}_\Omega) d\hat{\Gamma}_\Omega$ the characteristic function of '$P(\hat{\Gamma}_\Omega)$', by multiplying by '$e^{ik\hat{\Gamma}_\Omega}$' both right and left-hand sides of Eq.SM6.b-2 and integrating over any possible value of '$\hat{\Gamma}_\Omega$', we then deduce:

$$\widetilde{P}(k) = \int e^{ik\hat{\Gamma}_\Omega} \delta\left(\hat{\Gamma}_\Omega - F_\Omega(\{\Delta\hat{\theta}_j\})\right) \prod_{j=1}^{n} P_j(\Delta\hat{\theta}_j, \Omega) d\Delta\hat{\theta}_j \, d\hat{\Gamma}_\Omega$$

Or equivalently:

$$\widetilde{P}(k) = \int e^{ikF_\Omega(\{\Delta\hat{\theta}_j\})} \prod_{j=1}^{n} P_j(\Delta\hat{\theta}_j, \Omega) d\Delta\hat{\theta}_j \qquad \text{(SM6.b-3)}$$

As Eq.SM6.b-3 is a function '$F_\Omega$' and further assumptions are needed to provide its analytical form.

If we assume that the interaction between genome positions is primarily driven by the variations in the states of gene microstates involved, then we can assume that '$F_\Omega$' is independent of phenotype values and can thus be written more formally as 'F'. Therefore, the variable '$\hat{\Gamma}_\Omega$' will only result from the genome positions, irrespective of the environment. As the fields have been obtained considering the phenotype values and the phenotype results from the involvement of genome positions, any interactions between genome positions are already contained within the fields. As a result, the simplest formulation for 'F' is to consider a linear combination of $\Delta\hat{\theta}_j$'s such as:

$$F(\{\Delta\hat{\theta}_j\}) = \sum_j \Delta\hat{\theta}_j$$

Noting $\mu_j \stackrel{\text{def}}{=} \left(\Delta\hat{\Sigma}_1\right)_j(\Omega)$ and $\tau_j^2 \stackrel{\text{def}}{=} \left(\hat{\Sigma}_2\right)_j(\Omega)/N$ to simplify notations, using Eq.SM6-1 and Eq.SM6.b-3 we deduce:

$$\widetilde{P}(k) = \int e^{ik\Delta\hat{\theta}_j} \prod_{j=1}^{n} P_j(\Delta\hat{\theta}_j, \Omega) d\Delta\hat{\theta}_j$$

We then deduce by using Eq.SM6.b-1 that:

$$\widetilde{P}(k) = \prod_{j=1}^{n} \frac{1}{\sqrt{2\pi\tau_j^2}} \int e^{ikb_j\tau_j\left(\frac{\Delta\hat{\theta}_j}{\tau_j}\right) - \frac{1}{2}\left(\frac{\Delta\hat{\theta}_j}{\tau_j} - \frac{\mu_j}{\tau_j}\right)^2} d\Delta\hat{\theta}_j$$

Therefore,

$$\widetilde{P}(k) = \prod_{j=1}^{n} \frac{e^{ik\tau_j\left(\frac{\mu_j}{\tau_j}\right)-\frac{1}{2}\left(kb_j\tau_j\right)^2}}{\sqrt{2\pi}} \int e^{-\frac{1}{2}\left(\frac{\Delta\widehat{\theta}_j}{\tau_j}-\frac{\mu_j}{\tau_j}-ik\tau_j\right)^2} d\left(\frac{\Delta\widehat{\theta}_j}{\tau_j}\right) = \prod_{j=1}^{n} e^{-\frac{1}{2}\left[\left(k\tau_j\right)^2-2ik\tau_j\left(\frac{\mu_j}{\tau_j}\right)\right]}$$

Defining $\mu_{\widehat{\Gamma}_\Omega} \stackrel{\text{def}}{=} \left(\sum_{j=1}^{n} \mu_j\right)$ and $\tau^2_{\widehat{\Gamma}_\Omega} \stackrel{\text{def}}{=} \sum_{j=1}^{n}\left(\tau_j\right)^2$, the relation is then rewritten as:

$$\widetilde{P}(k) = e^{-\frac{1}{2}\left[\left(k\tau_{\widehat{\Gamma}_\Omega}\right)^2-2ik\mu_{\widehat{\Gamma}_\Omega}\right]}$$

By using the inverse Fourier transform of the characteristic function '$\widetilde{P}(k)$' one finds finally:

$$P\left(\widehat{\Gamma}_\Omega\right) = \frac{1}{2\pi}\int e^{-ik\widehat{\Gamma}_\Omega} e^{+ik\mu_{\widehat{\Gamma}_\Omega}-\frac{1}{2}k^2\tau^2_{\widehat{\Gamma}_\Omega}} d\widehat{\Gamma}_\Omega = \frac{1}{\sqrt{2\pi\tau^2_{\widehat{\Gamma}_\Omega}}} \exp\left[-\frac{1}{2}\left(\frac{\widehat{\Gamma}_\Omega-\mu_{\widehat{\Gamma}_\Omega}}{\tau_{\widehat{\Gamma}_\Omega}}\right)^2\right] \qquad \text{(SM6.b-4)}$$

Eq.SM6.b-4 is a version of the central limit theorem applied to the case of a set of genetic path differences. We deduce as expected that the average and variance of '$\widehat{\Gamma}_\Omega$' i.e $\langle\widehat{\Gamma}_\Omega\rangle = \int \widehat{\Gamma}_\Omega P\left(\widehat{\Gamma}_\Omega\right) d\widehat{\Gamma}_\Omega = \mu_{\widehat{\Gamma}_\Omega}$, and $\langle\left(\widehat{\Gamma}_\Omega\right)^2 - \langle\widehat{\Gamma}_\Omega\rangle^2\rangle = \tau^2_{\widehat{\Gamma}_\Omega}$ verify, respectively:

$$\mu_{\widehat{\Gamma}_\Omega} = \sum_{j=1}^{n} \mu_j = \sum_{j=1}^{n} \int_{\Omega_{1/N}}^{\Omega} \left[(\Delta\widehat{\omega})_j(\Omega) - (\Delta\omega_0)_j\right] \frac{1}{\Delta(\Omega)} d\Omega \qquad \text{(SM6.b-5)}$$

$$\tau^2_{\widehat{\Gamma}_\Omega} = \sum_{j=1}^{n}\left(\tau_j\right)^2 = \sum_{j=1}^{n}\left(\frac{1}{N}\int_{\Omega_{1/N}}^{\Omega}\left(1-(\widehat{\omega}_0)_j(\Omega)\right)\frac{1}{\Delta(\Omega)} d\Omega\right)^2$$

Eq.SM6.b-5 is then a generalized version of Eq.13 (main text).

**SM7: Signal-to-noise ratio when GIFT is used**

In the main text, we only considered the average value from the replicates to provide the best set of genotypes for a given population based on Fisher's theory.

However, as simulations were performed using $n = 1000$ replicates of $N = 1000$ individuals, it is possible with the simulated data to have an idea of the different signal-to-noise ratios obtained when the genetic path difference method is used.

As a number $n = 1000$ replicates were used, each representing $N = 1000$ individuals, it was possible for each replicate to rank the population of 1000 individuals as a function of the magnitude of phenotype values generated.

In this context, let us note '$\mathrm{g(i)}$' the average of genetic microstates found at the ranking position 'i'. The variable '$\mathrm{g(i)}$' can be rewritten as:

$$\mathrm{g(i)} = (+1) \times \frac{\mathrm{N_+(i)}}{n} + (0) \times \frac{\mathrm{N_0(i)}}{n} + (-1) \times \frac{\mathrm{N_-(i)}}{n}$$

where '$\mathrm{N_+(i)}$', '$\mathrm{N_0(i)}$' and '$\mathrm{N_-(i)}$' are, respectively, the number of microstates '+1', '0' and '-1' found at the position 'i' after performing $n = 1000$ replicates. Therefore, the ratios defined by '$\frac{\mathrm{N_+(i)}}{n}$', '$\frac{\mathrm{N_0(i)}}{n}$' and '$\frac{\mathrm{N_-(i)}}{n}$' correspond to the presence probabilities of finding, respectively, the genetic microstates '+1', '0' and '-1' at the position 'i'. Those presence probabilities can be assimilated to '$\omega_+(\mathrm{i})$', '$\omega_0(\mathrm{i})$' and '$\omega_-(\mathrm{i})$' as defined in the main text.

As a result,

$$\mathrm{g(i)} = (+1) \times \omega_+(\mathrm{i}) + (0) \times \omega_0(\mathrm{i}) + (-1) \times \omega_-(\mathrm{i}) = \omega_+(\mathrm{i}) - \omega_-(\mathrm{i})$$

Similarly, the variance of genetic microstates resulting from the replicates, '$\sigma_\mathrm{g}^2(\mathrm{i})$', is:

$$\sigma_\mathrm{g}^2(\mathrm{i}) = (+1)^2 \times \omega_+(\mathrm{i}) + (0)^2 \times \omega_0(\mathrm{i}) + (-1)^2 \times \omega_-(\mathrm{i}) = \omega_+(\mathrm{i}) + \omega_-(\mathrm{i})$$

Recalling Eq.SM6.a-1 one recognizes, therefore, that: $\Sigma_1 \sim \sum \mathrm{g(i)}$ and $\Sigma_2 \sim \sum \sigma_\mathrm{g}^2(\mathrm{i})$.

The signal-to-noise ratio can now be addressed as a function of the position when the genetic paths difference method is used. In this context, the argument in the exponential function of

Eq.SM6.a-1 provides a way to address this signal-to-noise ratio, noted '$\tau(\hat{\imath})$', for any position, as it is defined by:

$$\tau(\hat{\imath}) \stackrel{\text{def}}{=} \frac{\Sigma_1(\hat{\imath})}{\sqrt{\frac{2\Sigma_2(\hat{\imath})}{N}}}$$

In Figure SM7 (below), using simulated data, we provide the distribution of microstates for three log values of the gene effect, as well as the resulting '$\tau(\hat{\imath})$' for those gene effects. The conclusion reached is that the signal-to-noise ratio is relatively large (>1); therefore, GIFT is a method that can extract information efficiently. One way to compare the method of averages (Fisher's method) and GIFT is to compare Figure 1B (main text) with Figure SM7 because the same simulation parameters were used. Although it is difficult to extract a good signal-to-noise ratio with Fisher's method for small gene effects (Figure 1B), GIFT provides a robust method for extracting information with a restricted number of $N = 1000$ individuals. Note that Figure 3 (main text) suggests that with a gene effect of $a/\sigma = 0.01$, a study considering 1000 individuals would not allow 80% power to be achieved when the method of average is applied. In fact, Figure 3A demonstrates that with Fisher's method, the sample size would need to be approximately one million.

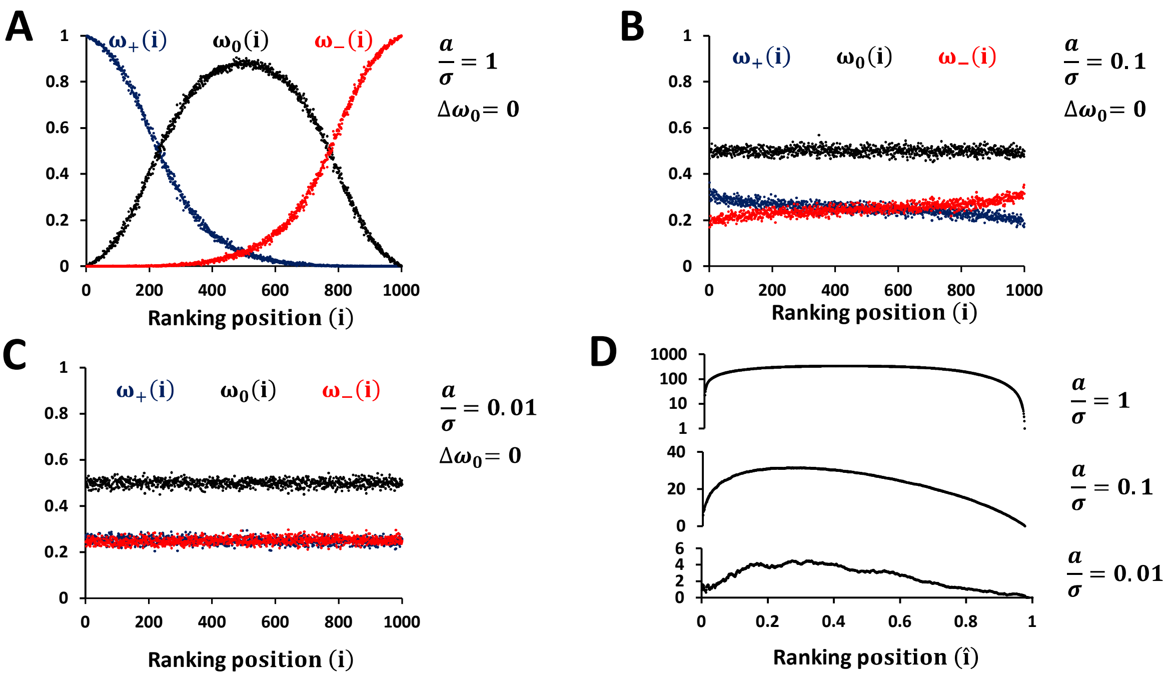


**Figure SM7: (A)** Representation of the presence probability of microstates as a function of positions obtained from the simulations when $a/\sigma = 1$, $\Delta\omega_0 = 0$ and $d = 0$. **(B)** Representation of the presence probability of microstates as a function of positions obtained from the simulations when $a/\sigma = 0.1$, $\Delta\omega_0 = 0$ and $d = 0$. **(C)** Representation of the presence probability of microstates as a function of positions obtained from the simulations when $a/\sigma = 0.01$, $\Delta\omega_0 = 0$ and $d = 0$. **(D)** Representation of the signal-to-noise ratios '$\tau(\hat{\imath})'s$' obtained from the genetic paths difference model using simulations. Note that the positions were normalised by the number of individuals in the population as explain in the main text. The conclusion is that the signal-to-noise ratio is higher than one for the log-value chosen of the gene effects.

**SM8: GIFT using real data**

GIFT relies on accurate phenotypic measurements. In this context, the 1001G+ project in plant (A. thaliana) is a fundamental resource for performing GWAS on a diverse range of plant phenotypes. The selected phenotype was the salt concentration in the plant leaves. Salt regulation in plants is an essential process regulating growth, and therefore, productivity as soil salinity and dehydration linked to climate change causes yield loss around the world (1). The high-affinity potassium transporter family (HKT1) has been found to be essential for reducing sodium toxicity linked to its accumulation in leaves (2), and genotype-phenotype associations/GWAS (3), as well as functional studies (4), have fully characterised the transporter. The advantages of using this system to demonstrate the relevance of GIFT are linked to the fact that sodium concentrations in leaves were determined with very high precision, but categories had to be artificially created to use GWAS (3). In Figure SM7, we present examples illustrating GIFT using this resource.

**SM7 references:**


1. Munns R, Tester M. Mechanisms of Salinity Tolerance. Annu. Rev. Plant Biol [Internet]. 2008 Apr 29;59(1):651–81. Available from: https://doi.org/10.1146/annurev.arplant.59.032607.092911
2. An D, Chen J-G, Gao Y-Q, Li X, Chao Z-F, Chen Z-R, et al. AtHKT1 drives the adaptation of Arabidopsis thaliana to salinity by reducing floral sodium content. PLoS Genet. 2017;13(10):e1007086.
3. Busoms S, Paajanen P, Marburger S, Bray S, Huang X-Y, Poschenrieder C, et al. Fluctuating selection of migrant adaptive sodium transporter alleles in coastal <em>Arabidopsis thaliana</em> Proc Natl Acad Sci [Internet]. 2018 Dec 26;115(52):E12443 LP-E12452. Available at: http://www.pnas.org/content/115/52/E12443.abstract
4. Uozumi N, Kim EJ, Rubio F, Yamaguchi T, Muto S, Tsuboi A, et al. The Arabidopsis HKT1 gene homologue mediates inward Na(+) currents in Xenopus laevis oocytes and Na(+) uptake in Saccharomyces cerevisiae. Plant Physiol. [Internet]. 2000 Apr;122(4):1249–59. Available from: https://pubmed.ncbi.nlm.nih.gov/10759522.

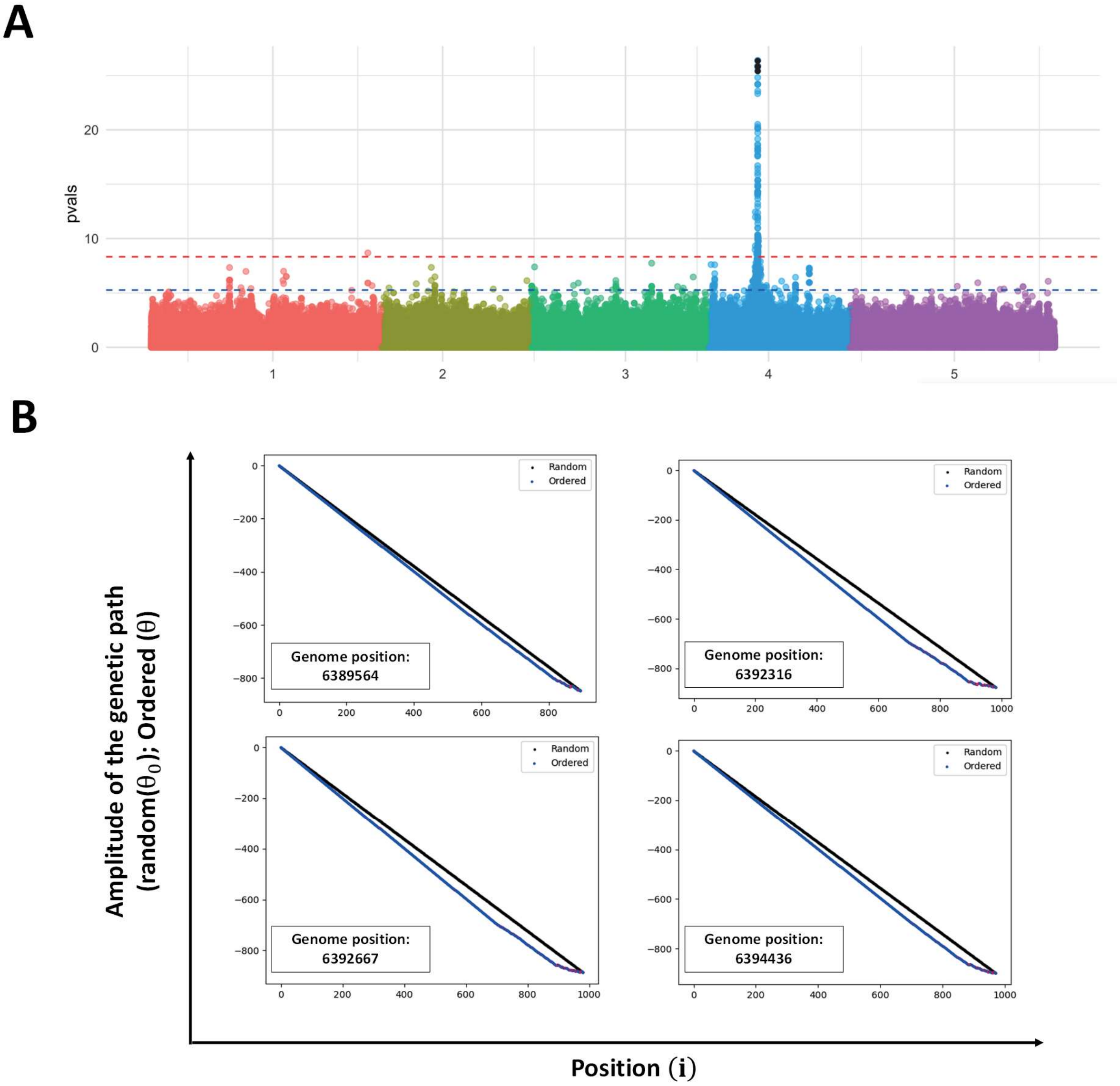


**Figure SM8**: Comparison of classic GWAS and GIFT using sodium leaf content. **(A)** Classic GWAS Manhattan plot. SNPs are coloured by chromosome (pink, yellow, green, blue and purple for chromosomes 1-5 respectively). The 'top 4' SNPs, i.e. the four most significant SNPs (lowest p-values), are shown in black at the top of the peak associated with HKT1, a sodium transporter experimentally shown to control leaf sodium content. Bonferroni threshold is shown as a red dashed line. Benjamini-Hochberg-Yekutieli threshold is shown as a blue dashed line. **(B)** Cumulative sum of genetic microstates for the 'top 4' classic GWAS SNPs (the genome positions are provided in the insets). The black line is the cumulative sum of genetic microstates upon the random ordering of phenotype values. The curved line is the cumulative sum of genetic microstates resulting from the ordering of phenotype values (in red are '+1' genetic microstates and in blue are '-1' genetic microstates). Note that we have used the GWAS convention to allocate the genetic microstates that is opposite to the one developed in the main text. Indeed, by convention GWAS allocates '+1' (resp. '-1') to genetic microstate characterising large (resp. small) phenotype values.
